# Lipoprotein lipase requires a flexible lid and stable C-terminal, both maintained by ApoC-II peptide binding

**DOI:** 10.1101/2025.11.20.689556

**Authors:** Emma E. Lietzke, Mary S. Rouse, Dean Oldham, Ziyue Dong, Robert H. Eckel, Kayla G. Sprenger, Kimberley D. Bruce

## Abstract

Lipoprotein lipase (LPL) is the rate-limiting enzyme responsible for hydrolyzing triglycerides in circulating lipoproteins. Reduced LPL activity contributes to hypertriglyceridemia, a major cardiovascular risk factor. LPL activity is thought to depend on the conformation of the lid domain, the lipid pore, N- and C-terminal domains (NTD, CTD), and stabilization of these domains by endogenous activators such as apolipoprotein C-II (ApoC-II). Despite major clinical significance, the structure-function relationship of LPL’s functional domains and cofactors remain incompletely understood. To address this, we performed the longest known (1-µs) molecular dynamics simulations of LPL independently and in complex with an ApoC-II mimetic peptide (ApoC-II-P). For the first time, we show that LPL’s flexible lid can adopt multiple orientations, transitioning between open and closed states that regulate lipid pore access and catalytic activity. We also observed ‘flipping’ of ∼180° by the CTD, a unique characteristic that dictates LPL activity when not in a closed lid state. Furthermore, ApoC-II-P stabilizes LPL by bridging its NTD and CTD, while maintaining an optimal lid orientation. Biochemical and cellular assays corroborate these findings, demonstrating that ApoC-II-P enhances LPL hydrolysis and supports noncanonical LPL functions. Together, these insights reveal previously unrecognized mechanisms governing LPL regulation and activity dynamics.

## Introduction

Lipoprotein lipase (LPL) regulates lipoprotein metabolism and lipid partitioning in key metabolic tissues such as the heart, adipose tissue, muscle^1,2^, and brain^3–5^. The canonical role of LPL involves the hydrolysis of triglycerides within triglyceride-rich lipoproteins, such as chylomicrons and very-low-density lipoproteins (VLDLs), liberating free fatty acids for cellular uptake^6,7^. Loss-of-function mutations in LPL have been repeatedly implicated in diseases, including familial hypertriglyceridemia^8^, cardiovascular disease^9^, and Alzheimer’s disease^10^. Such variants are thought to destabilize the functional domains of the enzyme, leading to loss of hydrolytic activity^7^. However, the precise mechanisms by which LPL’s structure dictates its activity remain an active area of investigation.

LPL contains numerous functional regions that mediate lipid, lipoprotein receptor, and cofactor binding. The N-terminal domain (NTD) includes an active site^11^ surrounded by a catalytic triad, oxyanion hole, and flexible lid domain that is thought to open and close, controlling lipid—and perhaps other ligand—accessibility to the site^12^ (Fig. 1A-B). Meanwhile, its C-terminal domain (CTD) contains a tryptophan-rich lipid-binding site (LBS)^13^ and a low-density lipoprotein receptor (-related protein) receptor binding site (LRP-BS)^14,15^, which also mediates self-association^16–18^. Like pancreatic lipase^19^, LPL was initially thought to only be active as a head-to-tail homodimer^20^, mediated by heparin binding. However, recent crystal structures reported helical head-to-tail oligomers^18^, tail-to-tail homodimers^21^, and active monomers and head-to-tail heterodimers bound to glycosylphosphatidylinositol-anchored high-density lipoprotein-binding protein 1 (GPIHBP1)^16,17,22^. In addition, the use of a small molecule as a stabilizing factor enabled crystallization of LPL with fully resolved lid and lipid-binding domains, revealing hydrophobic patches that allow for lipoprotein recognition^17^. Overall, these recent advances indicate more complex intra- and intermolecular interactions than previously recognized and galvanize the notion that LPL must be stabilized by ligands for proper functionality. However, our understanding of LPL dynamics in the presence or absence of such stabilizing ligands remains limited.

**Fig. 1:**
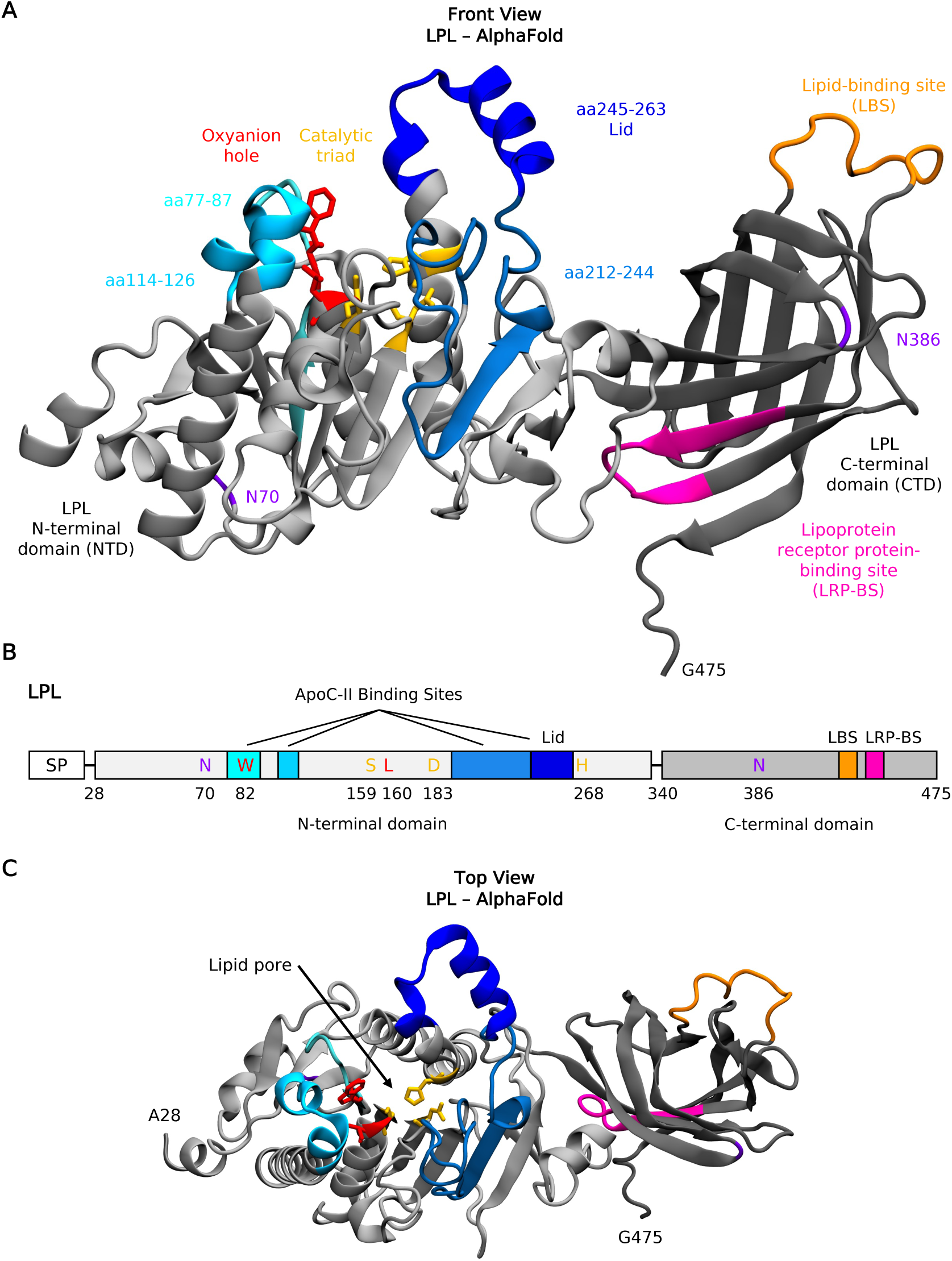
LPL contains multiple functional domains surrounding the ApoC-II binding sites. (A) The front view of the AlphaFold LPL monomer structure, numbered amino acids (aa) 28-475. LPL is shown with a cartoon representation. The N-terminal and C-terminal domains are shown in light and dark gray, respectively. At LPL’s catalytic site, the catalytic tried and oxyanion hole are shown with licorice representation, in yellow and red, respectively. The binding domains of ApoC-II are shown in cyan, sky blue, steel blue and dark blue, with their residue numbers listed. The lid domain is also labelled in dark blue. The lipid binding site and lipoprotein receptor-related protein (LRP) binding sites are shown in orange and pink, respectively. LPL’s N-linked glycosylation sites are shown in purple. (B) Schematic of LPL structure. All domains and residues listed in (A) are labelled. SP: signal peptide; LBS: lipid-binding site; LRP-BS: lipoprotein receptor protein-binding site. (C) The top view of LPL structure described in (A). The lipid pore is labeled, and the lid is seen in an open position, not covering the lipid pore.

Three small-molecule LPL activators have been identified, which can increase LPL expression or activity. However, they have limited applications in individuals carrying loss-of-function LPL polymorphisms or structural vulnerability^23–27^, as upregulating or targeting defective variants may not recover catalytic function. Given these limitations, therapeutics derived from endogenous LPL activators rather than exogenous compounds represent an attractive alternative strategy. Of these, Apolipoprotein C-II (ApoC-II) has been well studied. ApoC-II contains lipid and LPL binding sites on its N- and C-terminal helices, both thought to be essential for LPL function^28^. Historically, ApoC-II was thought to regulate LPL’s activity indirectly by altering lipid accessibility. However, recent studies have shown that ApoC-II actively regulates LPL’s catalytic activity. For example, mutagenesis assays identified individual residues within an LPL-activating region of ApoC-II that are necessary for LPL function^29^. In addition, a recent study found that ApoC-II activated monomeric LPL by binding to and anchoring the lid domain and interacting with and stabilizing the active site^30^, supporting the notion that effective activators stabilize LPL. Subsequently, a study discovered that a lipid pore spanning the NTD from top to bottom forms adjacent to the active site (Fig. 1C), exclusively in active LPL tail-to-tail homodimers^21^. However, it is unknown if ApoC-II binding directly impacts the opening and availability of the lipid pore and exactly how it mediates conformational changes in the lid. Molecular dynamics (MD) simulations offer a powerful approach to elucidate these atomic-level details that are difficult to assess experimentally. While molecular dynamics (MD) studies of LPL have been performed, previous studies used short timescales (e.g., 4 ns) and LPL structures without an intact lid^31–33^. Therefore, MD studies performed with intact structures and physiologically relevant timescales (e.g., within the µs range) is a desired approach to assess the molecular interaction between ApoC-II and LPL, and may guide the design of novel therapeutics.

We previously identified a mimetic ApoC-II peptide that significantly increases LPL activity^34^. Since Kumari et al.’s work identified that full-length ApoC-II could stabilize LPL^30^, we asked whether our peptide served a similar role. To do this, we performed 1-µs molecular dynamics simulations of LPL independently and LPL in complex with the ApoC-II peptide (ApoC-II-P). Interestingly, we discovered that LPL’s flexible lid can adopt multiple orientations, switching between open, intermediate, and closed states that are likely linked to active and inactive forms. Notably, we also observed CTD ‘flipping’ of ∼180° in the z-direction (with respect to an upright, stable NTD), which likely mediates LPL activity when it is not in an inactive, closed lid state. Outside these self-regulatory mechanisms, we found that the peptide can stabilize LPL by binding across its N- and C-terminal domains, thereby preventing CTD flipping. Experimental studies also revealed that ApoC-II-P increased the thermostability of LPL and increased LPL-mediated phagocytosis. Taken together, our findings provide new insights into the structure-function relationship between LPL and ApoC-II and support the need for further therapeutic development of ApoC-II peptides as LPL activators.

## Results

### LPL samples diverse lid orientations that control the accessibility of its lipid pore

Previous studies have identified numerous functional domains, but how these domains interact to influence function and stability remains unclear. To this end, we simulated monomeric LPL in aqueous solvent for 1 µs, in triplicate: the longest timescale LPL has been simulated with all-atom molecular dynamics (MD) to date. We first measured lid distance as a metric of lid opening. Importantly, our simulations recapitulated previous work^21^ showing that LPL’s lid sampled a dynamic range of conformations, resulting in open, intermediate, and closed states (Fig. 2A, see Fig. 2C for visualization).

**Fig. 2:**
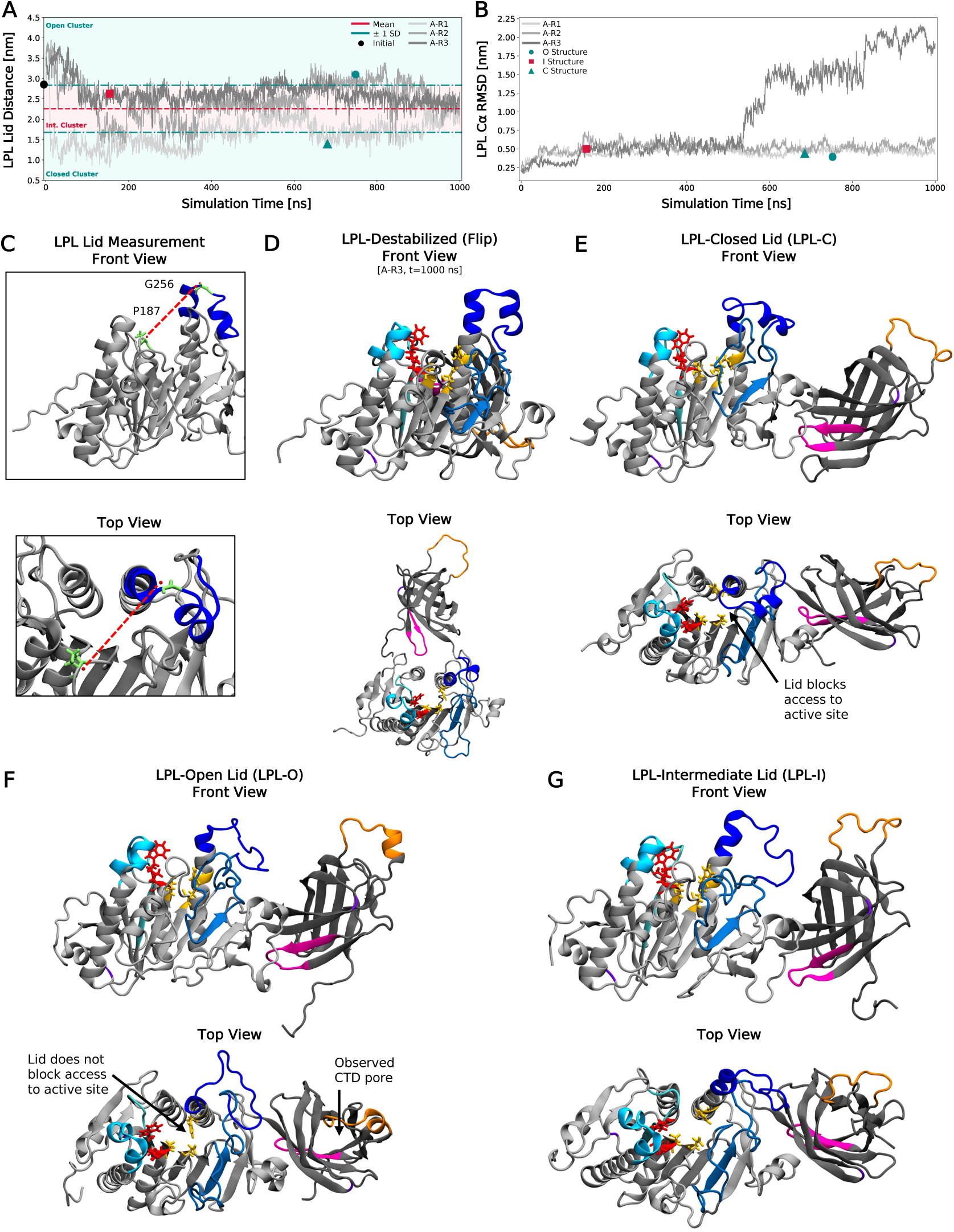
LPL samples various lid conformations when unbound to receptors, lipids, or cofactors. (A) LPL lid distance measured over time for the three LPL simulations. Lid distances for the replicates are plotted in gray. The average lid distance over the entire simulation time across all three simulations is plotted with a red, dashed line. The standard deviation (SD) above and below the mean are plotted with teal, dash-dotted lines. Lid distances above +1 SD are in the ”Open Lid Cluster”, distances below -1 SD are in the “Closed Lid Cluster,” and those in between the SD bounds are in the “Intermediate Lid Cluster.” The representative structure for each cluster are shown with colored points. (B) Root-mean-square deviation (RMSD) of the LPL Cα backbone plotted against simulation time. Replicates are shown in gray and representative cluster structures are shown in colored points. (C) Visualization of how lid distance was calculated between residues P187 and G256, seen in the front and top view. (D) Representative structure for the destabilized LPL structure (seen in A-R3 in Fig. 2B). (E) Representative structure for the Closed Lid Cluster (LPL-C). (F) Representative structure for the Open Lid Cluster (LPL-O). (G) Representative structure for the Intermediate Lid Cluster (LPL-I).

Without a binding partner, LPL experiences rapid time- and heat-associated loss in activity. We observed similar destabilization in one of our simulations (Alone-Replicate 3 (A-R3)), where the root-mean-square deviation (RMSD) of LPL sharply increases around 550 ns (Fig. 2B), primarily driven by changes in the C-terminal domain (CTD) of LPL. Specifically, the CTD flipped upside down, so that the lipid-binding site (LBS) and LRP-binding site (LRP-BS) swapped positions relative to the N-terminal domain (NTD). Furthermore, the CTD also moved away from its original position, shifting entirely behind the NTD (Fig. 2D, Front and Top Views). Going forward, we refer to this behavior as “CTD flipping”, which, in the absence of a binding partner, is stochastic.

Visual analysis of the simulation trajectories confirmed the lid positions and clusters quantified in Fig. 2A. These positions can be observed in the representative structures of each lid cluster. The closed, open, and intermediate lid LPL structures (LPL-C, LPL-O, LPL-I, resp.) are seen in Fig. 2E, 2F, and 2G, respectively (for A-R1-3 visualization, see Supplementary Fig. 1). We observed the LPL-C structure inhibiting access to the catalytic triad at the active site (Fig. 2E), which was accessible in the LPL-O and LPL-I states (Fig. 2F-G). Furthermore, when observing LPL from the top view, two pores were visible. The lipid pore in the NTD has been previously investigated^21^, and transports the fatty acyl chain products of triglyceride hydrolysis. We also noticed a second pore in the CTD of LPL, located below the lipid-binding site in all LPL structures. While the functionality of this pore is undetermined, it was nonetheless a notable artifact of the CTD conformation.

### CTD flipping limits the accessibility of LPL’s hydrolytic and lipid-binding domains

Next, we sought to characterize how CTD flipping, and structural differences in the lid region influenced changes in other functional domains. While the NTD was stable for the majority of the simulation (Fig. 3A, 200-1000 ns), the CTD was very unstable (Fig. 3B, 150+ ns in A-R2-3). Interestingly, denaturation of the CTD was not unique to the replicate that experienced CTD flipping. The regions of LPL that drive this instability were characterized with root-mean-square fluctuations of all LPL residues in the converged period of the simulations, from 900-1000 ns (Fig. 3C). Again, the NTD residues were largely stable, with slight fluctuations in the lid domain (dark blue region), while the CTD experienced much larger fluctuations (Fig. 3C). As expected, A-R3 experienced much larger fluctuations across the entire CTD, especially in residues within the LBS and those near the NTD (∼aa340-390): the pivot point of the flipping motion. Movement in the LBS was not unique to A-R3, and was the most dynamic region in the CTD, outside of the tail residues.

**Fig. 3:**
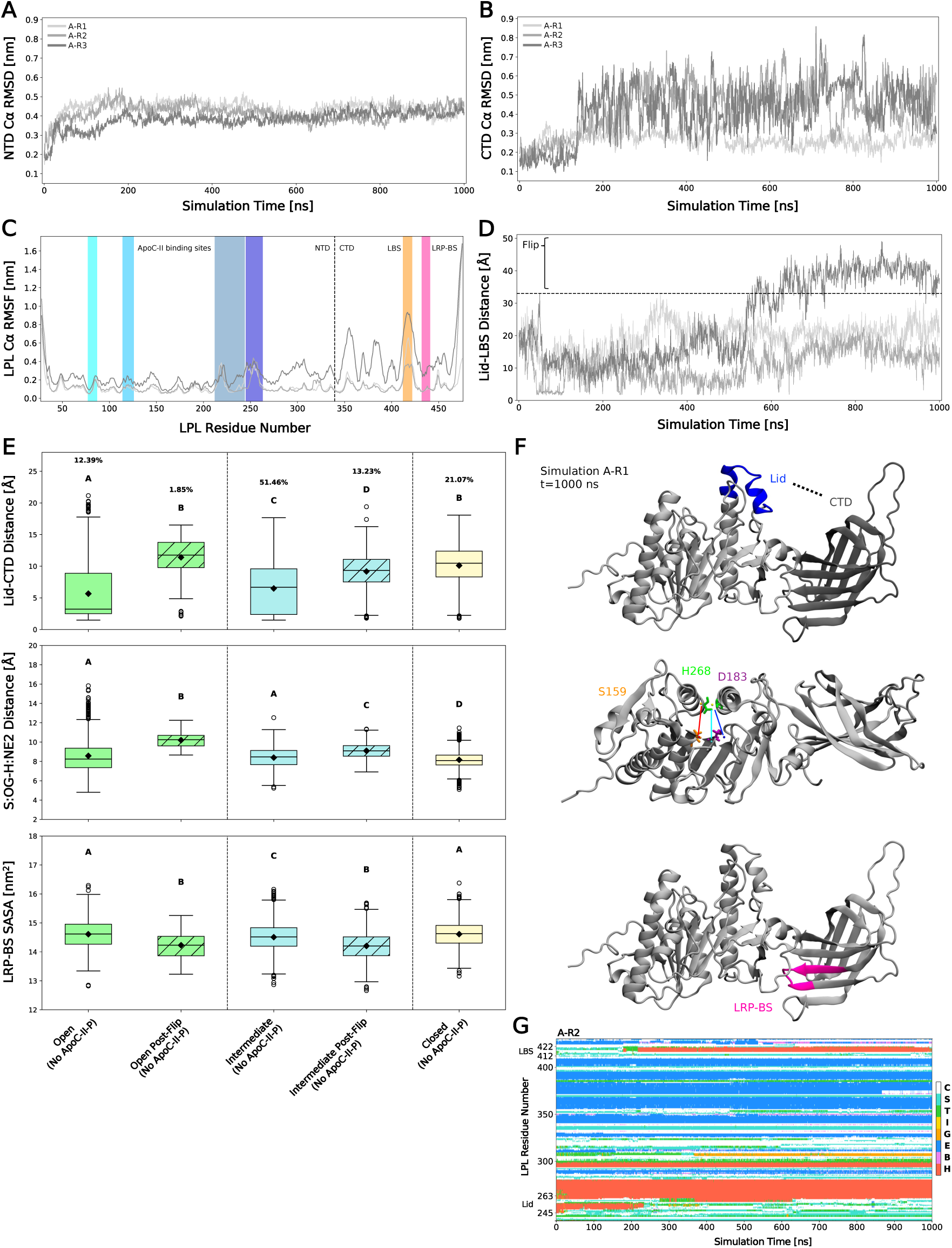
Functional domains of LPL are influenced by lid orientation and CTD flipping. (A) Root-mean-square deviation (RMSD) of the LPL N-terminal domain (NTD) Cα backbone plotted against simulation time. Replicates are shown in gray. (B) RMSD of the LPL C-terminal domain (CTD) Cα backbone plotted against simulation time. Replicates are shown in gray. (C) Root-mean-square fluctuation (RMSF) of the LPL Cα backbone over 900-1000 ns plotted against LPL residue number. Replicates are shown in gray. ApoC-II binding sites are shown in blue, the lipid binding site (LBS) in orange, and the receptor binding site (RBS) in pink. (D) Lid-LBS distance plotted against simulation time. Flipping is defined at a distance of 33 Å, and any points after this distance are classified as post-flipping. (E) Open lids are shown in green, intermediate in blue, and closed in yellow. Post-flip groups shown with hatching. Group occupation percentages, over total simulation time, are shown for each group. Compact-letter-display represents significantly different groups from a Kruskal-Wallis ANOVA with post-hoc Dunn’s comparison tests (N=3 independent simulation runs, with 2,001 data points/run; see Supplementary Table 1 for n breakdown per group). Top: lid-CTD distance measured across groups; middle: distance between the S159:OG and H268:NE2 atoms across groups; bottom: solvent-accessible surface area (SASA) of the LRP-BS across groups. (F) Visualization of the quantified domain metrics of LPL in (E). (G) Dictionary of protein secondary structure (DSSP) assignments for A-R2 per LPL residue number plotted against time. Assignments codes are: α-helix (H), residue in isolated β-bridge (B), extended strand participating in β-ladder, 3/10-helix (G), π-helix (I), hydrogen-bonded turn (T), bend (S), and loop/irregular elements (C)

Having established the temporal and spatial nature of CTD flipping, we sought to create a metric that quantified this motion. To do so, the minimum distance between residues in the lid (aa245-263) and the LBS (aa412-422) was calculated over the entire simulation period. Indeed, the Lid-LBS distance (Fig. 3D) of A-R3 was higher than that observed in A-R1 and A-R2. Given that flipping is likely to cause irreversible LPL inactivation, we sought to cluster LPL structures into pre- and post-flip groups, and set a threshold to determine the flipping point at 33 Å (Fig. 3D, black dashed line). After this distance, structures were considered “Post-Flip” (Fig. 3E). Structures that never sampled a flip were clustered together as a separate group, with no notation of a flip in the name (Fig. 3E).

Next, we postulated that variable lid states may drive functional changes that facilitate LPL-mediated hydrolysis, receptor binding^14–16,35^, and ligand bridging^14,15,36,37^. When measuring the distance between the lid and CTD, we found that lid orientation and flipping impacts the Lid-CTD distance (Fig. 3E, top; Fig. 3F, top, for visualization). As expected, a closed lid increased the Lid-CTD distance compared to intermediate and open-lid structures (Fig. 3E, top). Flipping also increased Lid-CTD distances in structures with open and intermediate lids, with flipped distances being comparable to that of a closed lid (Fig. 3E, top). Further, flipping resulted in greater distances between the lid and the LBS (Supplementary Fig. 2A). A shorter Lid-CTD distance in the open lid structures aligns with the lid’s potential to stabilize the CTD when it is not receptor-bound. Open lids were largely sampled by simulation A-R2, which exhibits consistent H-bonding between the lid and CTD throughout the simulation period (Supplementary Fig. 3A). This suggests that LPL’s lid might have an additional role in maintaining structural integrity outside of mediating pore accessibility and hydrolytic activity.

Secondly, we measured the distance of the donor-acceptor pairs in residues involved in the catalytic triad near the top of the lipid pore. We again found that flipping increased the distance between the O and N atoms in the S159-H268 H-bond (Fig. 3E, middle). Remarkably, we observed few significant differences despite lid orientation or flipping in the potential bonds between D183 and H268 (Supplementary Fig. 2B). This may suggest that the D183-H268 bond is relatively stable despite LPL status, while S159-H268 bonding is critical for stable catalytic activity and can be lost easily. In contrast, when we measured the radius of gyration of the pore-lining residues, we found that this region had similar compactness across open, closed, and open post-flip states (Supplementary Fig. 2C), illustrating that the lipid pore structure remains consistent across group types.

Since LPL is known to bind to a multitude of receptors, or even itself at its LRP-BS, we characterized the availability of this region across groups. We found that flipping decreased the solvent-available surface area (SASA) of the LRP-BS (Fig. 3E, bottom), further supporting the notion that flipping has significant impacts on LPL structure, function, and binding interactions at this region.

Finally, we characterized the dictionary of protein secondary structure assignments (DSSP) of LPL over time to quantify structural changes. Indeed, in A-R2, we observed a loss of α-helicity in the lid domain, which likely allows lid flexibility and interactions with the CTD and LBS (Fig. 3G). Intriguingly, when the lid loses helicity, helicity is gained in the LBS in A-R2. This behavior is not observed in A-R1 and A-R3, although these models do not sample an open lid conformation (see Supplementary Fig. 4 for all residues). To test whether helicity in the LBS leads to lipid binding, we docked three lipid species (oleic acid, phosphatidylserine, phosphatidylcholine) to the LBS of the LPL-O (helical LBS) and LPL-C (no helicity) structures (Supplementary Fig. 5A). Notably, we observed oleic acid (C18:1) preferentially binds to a helical LBS. Whether this relates to LPL’s preference for monounsaturated fatty acids remains to be determined^5,38^. Phospholipids did not preferentially bind to a helical LBS (Supplementary Fig. 5B), suggesting that different LPL conformations drive lipid preference. Overall, a helical LBS, paired with an open lid and accessible pore, may represent an optimal structure for LPL activity. Since we observe these structural phenomena in soluble, unbound LPL, next we probed whether the results would be recapitulated in the presence of a potentially stabilizing ligand: ApoC-II peptide (ApoC-II-P)^34^.

### The secondary structure of the ApoC-II peptide is dynamic

We previously designed an ApoC-II peptide (ApoC-II-P), which contains residues S78-E100 comprising ApoC-II’s LPL activating region, and profoundly activates LPL in triglyceride hydrolysis assays^34^. In order to accurately model LPL-ApoC-II-P interactions, we first sought to simulate ApoC-II-P independently for 1 µs, in triplicate (see Fig. 4A-B for peptide visualization). Peptide simulations showed increased RMSD over time across all replicates (Fig. 4C), indicating instability in the initial structure and thus large structural changes. This is supported by trajectory snapshots (Fig. 4F), which confirm the peptide consistently loses α-helicity over time. Large spikes in RMSD were compared to their respective snapshots, further validating that RMSD spikes correlate with large losses in secondary structure motifs (e.g., 200-400 ns for P-R2, 200 ns for P-R3). Indeed, the post-simulation representation structure of ApoC-II-P exhibits loss of helicity (Fig. 4D-E), with residues more likely to engage in β-sheets formation compared to the initial structure (Fig. 4A,E). This propensity for β-sheet formation aligns with prior reports of lipid-free full-length and peptide portions of ApoC-II forming morphologically variable amyloid fibrils^39–41^.

**Fig. 4:**
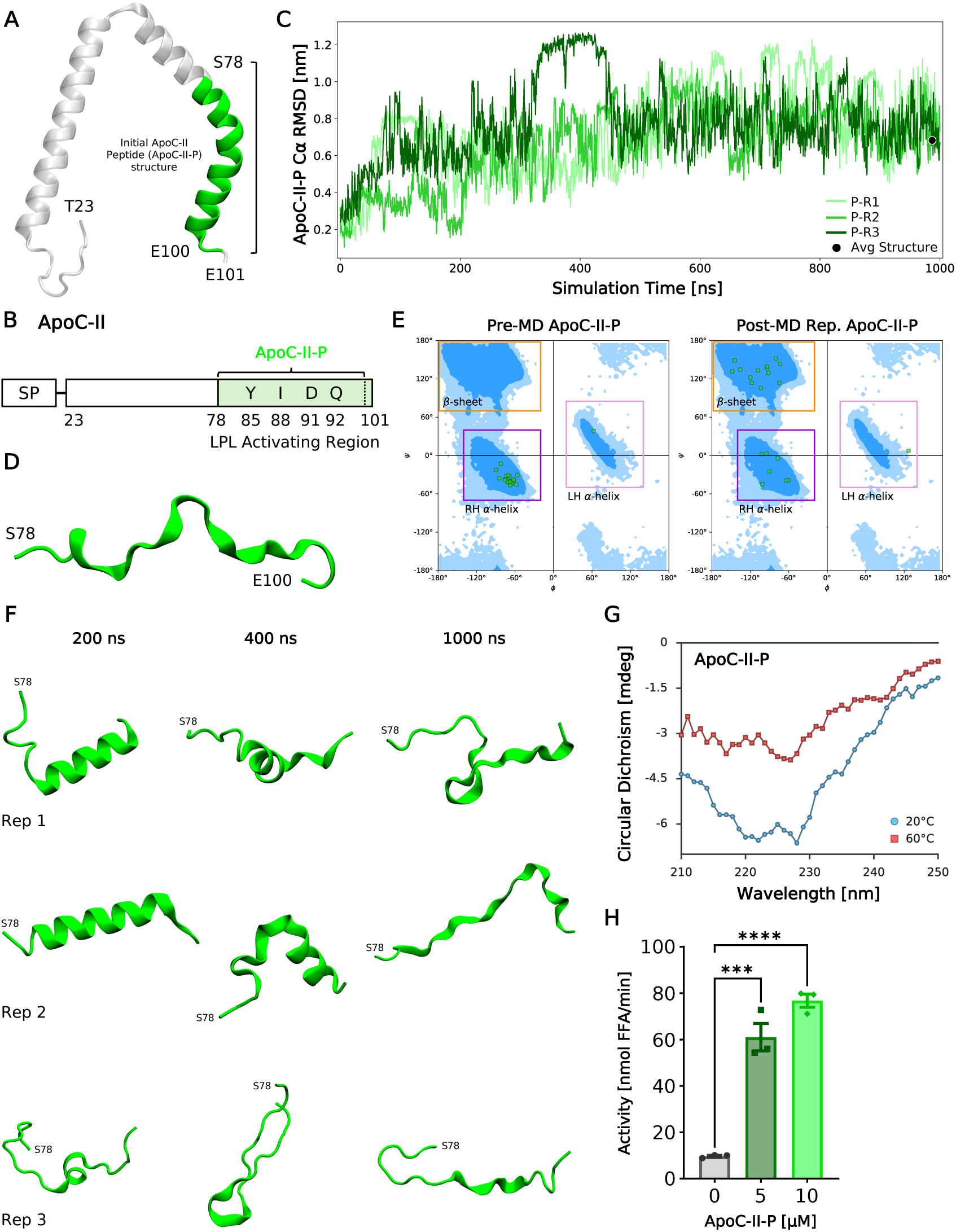
ApoC-II peptide (ApoC-II-P) loses α-helicity in simulations. (A) ApoC-II structure shown in cartoon. Full ApoC-II is shown in white, transparent cartoon. ApoC-II-P is shown in green, overlayed on ApoC-II, where it has been clipped from the full protein. (B) Root-mean-square deviation (RMSD) of the ApoC-II-P Cα backbone plotted against simulation time. Replicates are shown in green and representative cluster structures are shown in the black point. (C) The post-MD representative structure of ApoC-II-P from clustering the last 100 ns of each simulation. ApoC-II-P is shown in the same representation as in (A). (D) Ramachandran plots to describe secondary structure motifs of ApoC-II-P of the pre-MD structure (left; seen in Fig. 4A) and the post-MD representative (rep.) structure (right; seen in Fig. 4C). (E) Schematic of ApoC-II structure with key LPL binding residues listed. SP: signal peptide. (F) Trajectory snapshots of ApoC-II-P across the three replicates over simulation time. (G) Circular dichroism spectra of ApoC-II-P from 210-250 nm collected at 20°C and 60°C (n=1). (H) Activation of bovine LPL with increasing ApoC-II-P concentration (n=3). Student’s t-test was used for significance (*** = p<0.001, **** = p<0.0001).

These structural changes were further verified through circular dichroism (CD) experiments. ApoC-II-P exhibited weak α-helical character at 20°C, indicated by a modest negative ellipticity near 225 nm (Fig. 4G), but overall lacked defined secondary structure. Upon incubation above room temperature, ApoC-II-P completely lost residual structure, as reflected by the flattened, less negative spectra at 225 nm (Fig. 4G). These observations are consistent with our simulations, which show that ApoC-II-P progressively loses secondary structure over time. Next, we set out to confirm that ApoC-II-P could increase LPL activity despite its apparent loss in secondary structure. Indeed, we observed that the ApoC-II-P could profoundly increase hydrolytic activity in a dose-dependent manner (Fig. 4H), consistent with previous results^34^. Thus, we sought to characterize, for the first time, the mechanism by which unstructured ApoC-II-P increased and maintained the hydrolytic activity of LPL, by modeling these interactions with MD simulations and biochemical assays.

### LPL retains lid flexibility when bound to the ApoC-II peptide

Post-MD representative lid structures of LPL (-open(O), -intermediate(I), -closed(C)) were docked with a representative ApoC-II-P structure. Models were assessed for biological relevance by measuring the number of LPL residues at experimentally determined ApoC-II binding sites^30^. Top-scoring LPL-ApoC-II-P models, one for each of the three LPL orientations, were simulated for 1 µs, in triplicate, resulting in a total of nine simulations (denoted with a docking position dependent on lid orientation (X, Y, Z) and replicate number (R1, R2, R3); see Table 1 in **Methods** for metric and cluster descriptions, and Supplementary Fig. 6 for docking poses).

**Table 1.** Simulation set, cluster, and replicate descriptions.

| Simulation Set or Cluster | Replicates (R) | Description |
| --- | --- | --- |
| Set A | A-R1, A-R2, A-R3 | Technical replicates of LPL-Alone simulations, later referred to as No ApoC-II-P simulations of LPL. LPL simulated as a monomer in aqueous solvent. Output and clustering of simulations resulted in LPL-Open (O), LPL-Intermediate (I), and LPL-Closed (C) lid structures. |
| Set P | P-R1, P-R2, P-R3 | Technical replicates of the ApoC-II-P simulations. ApoC-II-P simulated, alone, in aqueous solvent. Output and clustering of simulations resulted in the |
|  |  | post-MD representative ApoC-II-P structure. |
| Cluster X | X-R1, X-R2, X-R3 | Technical replicates of simulations with the LPL-Open (LPL-O) lid structure docked to the rep. ApoC-II-P structure. Availability of LPL residues upon ApoC-II-P docking were dependent on an **open** lid. |
| Cluster Y | Y-R1, Y-R2, Y-R3 | Technical replicates of simulations with the LPL-Intermediate (LPL-I) lid structure docked to the rep. ApoC-II-P structure. Availability of LPL residues upon ApoC-II-P docking were dependent on an **intermediate** lid. |
| Cluster Z | Z-R1, Z-R2, Z-R3 | Technical replicates of simulations with the LPL-Closed (LPL-C) lid structure docked to the rep. ApoC-II-P structure. Availability of LPL residues upon ApoC-II-P docking were dependent on a **closed** lid. |

We observed unstable LPL in several simulations despite the binding of ApoC-II-P. RMSD of the complex was measured over time (Fig. 5A), with three replicates (Y-R1, Y-R3, Z-R3) showing significant RMSD spikes (CTD flipping), one replicate (X-R1) exhibiting a steep spike and then drop-off in RMSD (ApoC-II-P dissociation and reassociation), and others showing a gradual increase over time. RMSF of LPL in the converged period (900-1000 ns) was measured to assess the stability of the NTD and CTD regions (Fig. 5B). We observed the highest fluctuations in LPL across the CTD at the hinging point and LBS, but also high variability in the surrounding lid domain residues. As expected, the greatest LPL flexibility was observed in replicates with high RMSD profiles.

**Fig. 5:**
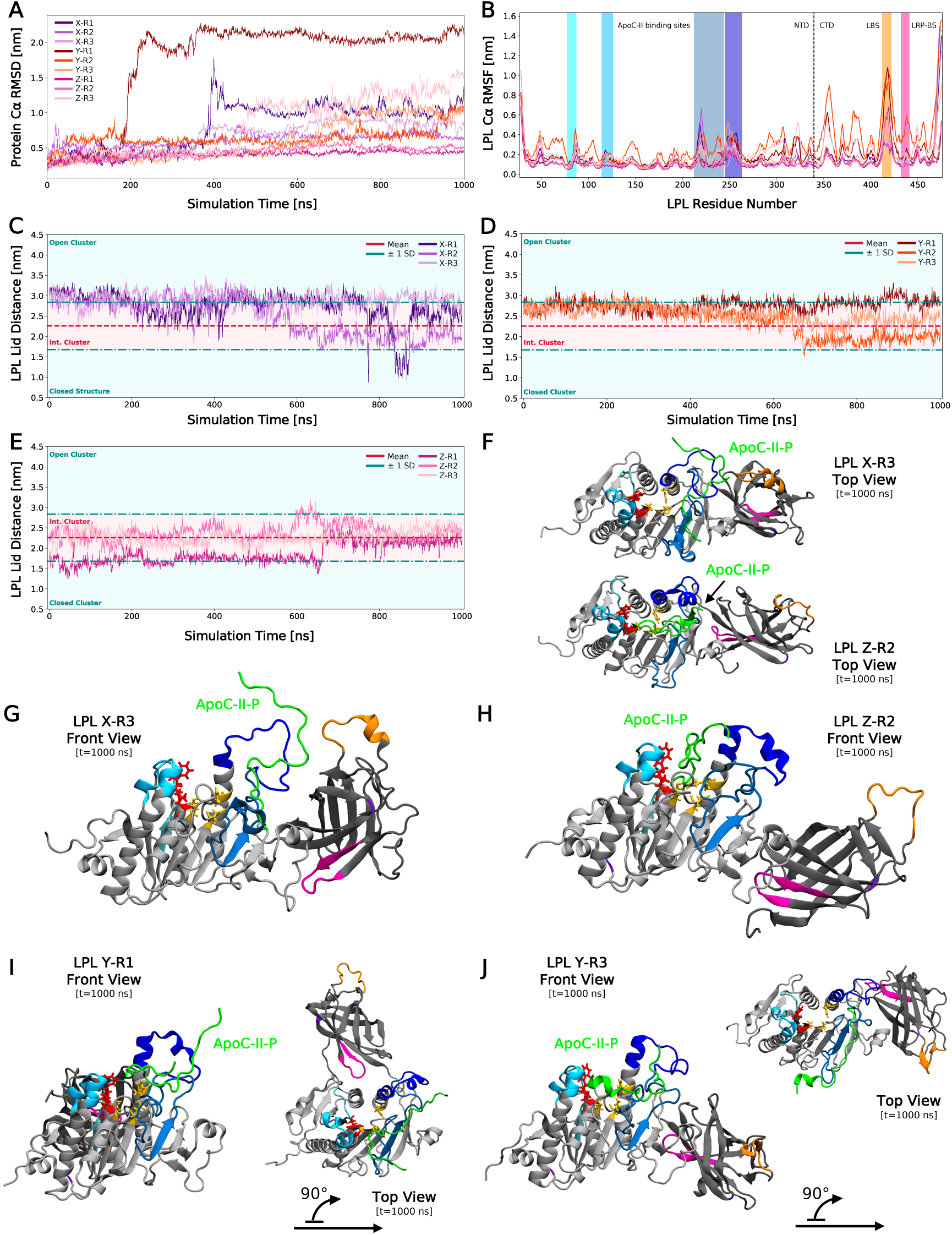
ApoC-II-P does not stabilize the LPL lid. (A) Root-mean-square deviation (RMSD) of the LPL Cα backbone plotted against simulation time. Replicate colors are based on the initial starting position from docking. Cluster X replicates are shown in purple shades, Y in orange shades, and Z in pink shades. (B) Root-mean-square fluctuation (RMSF) of the LPL Cα backbone over 900-1000 ns plotted against LPL residue number. Labels are listed and colored as described in Fig. 3C. (C) LPL lid distance of Cluster X replicates measured over time. Labels and colors for distance clusters are the same as those described in Fig. 2A. (D) LPL lid distance of Cluster Y replicates measured over time. (E) LPL lid distance of Cluster Z replicates measured over time. (F) Final time-point LPL visualization, in the top view, of X-R3 (top) and Z-R2 (bottom) with ApoC-II-P (green). LPL colors are those described in Fig. 1A. (G) Final time-point visualization of X-R3. (H) Final time-point visualization of Z-R2. (I) Final time-point visualization of Y-R1. (J) Final time-point visualization of Y-R3.

We then asked whether lid distance was stabilized by ApoC-II-P. Lid distance across all nine simulations were plotted over time, with the same clustering bounds identified in Fig. 2A. Surprisingly, ApoC-II-P binding did not stabilize lid orientation, especially in Cluster X simulations (Fig. 5C). Lid orientation was more stable in Clusters Y and Z upon peptide binding (Fig. 5D-E). Additionally, because the RMSD spikes and thus CTD flipping occurred at similar time points, we measured the lid distance at the beginning of this motion. For Y-R1, Y-R3 and Z-R3, flipping begins at 196.5, 713.5, and 470 ns, respectively. All distances at these time points are within the intermediate lid cluster, supporting the notion that a specific lid orientation is not required for CTD flipping. Conversely, an intermediate lid was sampled more frequently in the presence of ApoC-II-P (75%), suggesting for the first time that ApoC-II-P maintains a flexible lid orientation consistent with enzyme activity.

### ApoC-II peptide stabilizes LPL through bridging its N- and C-termini

Since previous studies have suggested that ApoC-II stabilizes LPL by anchoring the lid^29,30^, we next probed the mechanism of LPL activation given the maintenance of a flexible lid domain in the presence of ApoC-II-P. Notably, we observed ApoC-II-P binding at the top of the pore in Cluster Z simulations, potentially inhibiting lipid access to this region. Differential ApoC-II-P binding sites across Clusters X, Y, and Z are seen in Fig. 5F-J (see Supplementary Fig. 7 for all poses). Notably, ApoC-II-P can bridge the NTD and CTD (Fig. 5F-G). Overall, the orientation of the LPL lid significantly impacts where ApoC-II-P can bind, and thus its ability to bridge the C-and N-termini and potentially stabilize LPL and maintain catalytic activity.

Building on the idea that the ApoC-II-P binding location promotes LPL activity, we analyzed the impact of ApoC-II-P binding on LPL’s functional domains. Again, we observed that the NTD of LPL was significantly more stable than the CTD, independent of ApoC-II-P binding or CTD flipping (Fig. 6A-B). The dynamic nature of the CTD was further illustrated by quantification of CTD flipping and Lid-LBS measurements across the nine simulations (Fig. 6C). Three simulations experienced flipping (Y-R1, Y-R3, Z-R3) while those of Cluster X exhibited no flipping, with the NTD and CTD moving closer together (e.g., lid and LBS). Intriguingly, when comparing the No ApoC-II and with ApoC-II-P Lid-LBS distances, we observe that No ApoC-II simulations behave similarly to Cluster Y simulations. This observation suggests that LPL’s lid position, upon initial contact with ApoC-II-P, significantly impacts the peptide’s ability to stabilize LPL after initial binding.

**Fig. 6:**
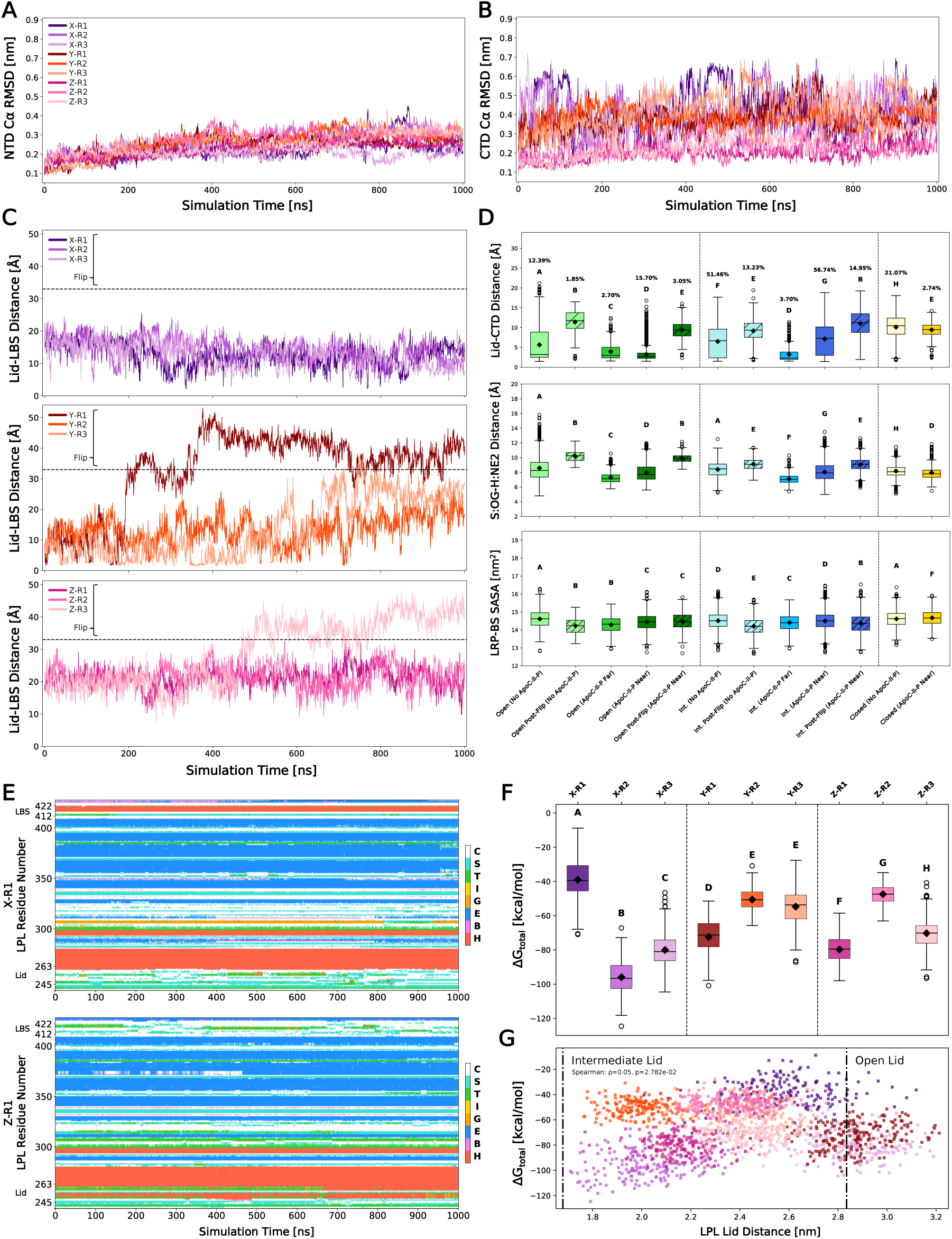
ApoC-II-P does not significantly affect LPL functional domains but binds differently to LPL based on initial lid orientation. (A) Root-mean-square deviation (RMSD) of the LPL N-terminal domain (NTD) Cα backbone plotted against simulation time. Replicate colors are the same as those described in Fig. 5A. (B) RMSD of the LPL C-terminal domain (CTD) Cα backbone plotted against simulation time. (C) Lid-LBS distance plotted against simulation time. Flipping is defined at a distance of 33 Å, and any points after this distance are classified as post-flipping. (D) Open lids are shown in green, intermediate in blue, and closed in yellow. Post-flip groups shown with hatching. Group occupation percentages, over total simulation time, are shown for each group. Groups represented less than 1% of the simulation time are not shown. Compact-letter-display represents significantly different groups from a Kruskal-Wallis ANOVA with post-hoc Dunn’s comparison tests (N=12 independent simulation runs, with 2,001 data points/run; see Supplementary Table 1 for n breakdown per group). Top: lid-CTD distance measured across groups; middle: distance between the S159:OG and H268:NE2 across groups; bottom: solvent-accessible surface area (SASA) of the LRP-BS across groups. (E) Dictionary of protein secondary structure (DSSP) assignments for X-R1 and Z-R1 per LPL residue number plotted against time. Assignments codes are: α-helix (H), residue in isolated β-bridge (B), extended strand participating in β-ladder, 3/10-helix (G), π-helix (I), hydrogen-bonded turn (T), bend (S), and loop/irregular elements (C). (F) Binding interaction energy between LPL and ApoC-II-P for 900-1000 ns (N=9 independent simulation runs, with 201 data points/run) was determined as ΔG_total_ and plotted for each simulation run. CLD represents significantly different groups from a Kruskal-Wallis ANOVA with post-hoc Dunn’s comparison tests. (G) ΔG_total_ values for each simulation run were plotted as a function of LPL lid distance. Dash-dot lines represent the boundaries for the closed, intermediate, and open lids. A Spearman’s test was performed to determine potential monotonic correlation. Colors represent the same groups as seen in (A-C, F) and the same N values as (F).

Next, we probed whether the presence of ApoC-II-P altered the impact of flipping on LPL’s functional domains. We again observed that CTD flipping significantly increased the distance between the lid and CTD regardless of lid orientation (Fig. 6D, top; for Lid-LBS measurements, see Supplementary Fig. 8A). This aligns with our observation that Cluster X, where ApoC-II-P tethered the C- and NTDs, helped to prevent flipping rather than stabilize the lid. Interestingly, intermediate and open lids were sampled after flipping events only when ApoC-II-P was close to its defined binding sites^30^, and a closed lid was never sampled by a flipped LPL complex. This further supports the notion that ApoC-II-P maintains lid flexibility while also preventing a closed lid state, promoting enzyme activity.

The trends observed in the catalytic triad and LRP binding site (LRP-BS), due to flipping, also occurred in the presence of ApoC-II-P. CTD flipping again significantly decreased the propensity for H-bonding at the catalytic triad, specifically between S159 and H268 (Fig. 6D, middle). In most cases, the availability of the LRP-BS also decreased when flipping occurred (Fig. 6D, bottom). Similarly, the presence of ApoC-II-P did not alter stable regions unimpacted by lid orientation or flipping (Supplementary Fig. 8B-D). Overall, this suggests that ApoC-II-P does not affect the structure of the catalytic triad and LRP-BS, but rather stabilizes LPL by bridging the N-and C-terminal domains.

Indeed, N-CTD bridging in Cluster X was further observed in the increased number of H-bonds between the lid and CTD (Supplementary Fig. 3B). This aligns with the secondary structure results, suggesting that an open lid, with the ability to bridge these domains, may be linked to increased α-helicity in the LBS rather than LBS’s interaction with the remaining CTD (Fig. 6E; see Supplementary Figs. 9-11 for full profiles).

Based on our hypothesis that ApoC-II-P forms a stabilizing bridge between the NTD and CTD, we measured the interaction energies (Fig. 6F) between LPL and ApoC-II-P over the converged, last 100 ns of the simulations. Considering that lid orientation and flipping impacted other functional domains, we wondered if CTD flipping or lid distance resulted in very weak or strong binding. However, the three simulations with flipped domains did not illustrate the weakest nor strongest binding (Fig. 6F), and continuous lid distance was not correlated to binding energy (Fig. 6G). Instead, we found that the weakest interaction energy occurred in one replicate of Cluster X when ApoC-II-P dissociated before convergence and re-associated at the LRP-BS, which is not ApoC-II’s primary binding site^30^. The strongest energies occurred in the remaining two Cluster X simulations, where ApoC-II-P bridged the C and N-termini (Fig. 6F), further highlighting the importance of this specific interaction.

Since lid distance and CTD flipping did not dictate strong binding, we sought to identify the residues mediating these interactions. To this end, we calculated the contact frequency across all LPL residues in the converged time (Supplementary Fig. 12; see Supplementary Table 2 for residue list) and paired it with a decomposition analysis (Supplementary Figs. 13-14) to identify residues with the strongest contributions. As anticipated, simulations X-R2 and X-R3 showed high likelihood of contact with CTD, and X-R1 showed the fewest interactions overall (Supplementary Fig. 12A). Contact frequencies generally aligned with binding energies, as simulations with a higher number of contacts, especially those in the defined ApoC-II-P regions^30^ (Supplementary Fig. 12B), had stronger binding.

Decomposition analyses further showed that the most frequent and strongest contributions were from R214 and R219, which reside in LPL’s lid adjacent region near the lipid pore (Supplementary Fig. 13A-B). While other lid residues were strong contributors, they did not occur as frequently, further suggesting that ApoC-II-P binding is anchored by peripheral pore regions rather than solely by the lid. This aligns with our observation that the lid remains dynamic, preferring an intermediate and open state compared to a closed conformation, likely due to nearby interactions. Finally, we identified that D91 and Q92 were the most common energetic contributing residues in ApoC-II-P, but these contributions were significantly lower when compared to the strength of the LPL residues (Supplementary Fig. 14A-B). This further implies that the length and composition of ApoC-II-P allow it to bind broadly to LPL. Still, the accessibility of LPL binding interfaces dictates where the peptide will actually bind. Therefore, dynamic protein characteristics such as lid position ultimately drive which interfaces become ideal binding spots to enable its stabilizing effect. This also suggests that while ApoC-II-P may maintain an active configuration of LPL, it may not be sufficient to rescue LPL activity when already in an inactive state.

### ApoC-II-P mitigates heat-induced inactivation of LPL and promotes LPL-mediated phagocytosis

Based on our findings that ApoC-II-P significantly impacted LPL structure depending on its initial contact position, we wondered if this would be reflected in CD spectra of LPL with and without ApoC-II-P. We observed a modest difference in the 220-240 nm range when ApoC-II-P was present (Fig. 7A). This might indicate that ApoC-II-P induces a slight alteration in the secondary structure of LPL via stabilization or local rearrangement, aligning with our MD results that ApoC-II-P can have stabilizing effects between LPL’s NTD and CTD. However, due to the loss of a positive peak at 190 nm, and less negative signals in the 208 and 225 range, these results may reflect the heterogeneity of LPL+ApoC-II-P structures and the propensity of LPL to access variable states when in complex with ApoC-II-P, consistent with our results from the MD simulations. Outside of structural changes, we wanted to verify that ApoC-II-P stabilizes and preserves the hydrolytic activity of LPL by performing radiometric bovine LPL (bLPL) activity assays, as previously described^5,34^. Specifically, to determine whether ApoC-II-P could prevent heat-mediated destabilization of LPL, we quantified LPL-mediated hydrolysis, when the optimal hydrolysis reaction (45 mins at 37°C) was preceded by a 30-minute incubation period at different temperatures (20°C, 40°C, and 60°C) with or without ApoC-II-P (Fig. 7B). Initially, ApoC-II-P was either present or absent during the pre-incubation period and was also present in the LPL substrate. LPL activity was highest in samples that were incubated at 20°C, with the samples treated with ApoC-II-P showing the highest activity levels, but not significantly (Fig. 7C). LPL activity was lower in both + and – ApoC-II-P conditions pre-incubated at 40°C. However, the loss of LPL activity at 40°C was not as pronounced in samples incubated with ApoC-II-P. Indeed, at 40°C, LPL activity was significantly higher in the presence of ApoC-II-P (Fig. 7C). At 60°C, all hydrolytic activity was lost, with or without the addition ApoC-II-P (Fig. 7C). Overall, this data suggests ApoC-II-P may increase the thermostability of LPL. However, the modest magnitude of an effect suggests that the addition of ApoC-II-P to the LPL substrate may have partially rescued the hydrolytic activity of LPL, by potentially stabilizing any enzyme that had not yet been irreversibly denatured, especially at 20°C. This notion is supported by the absence of an effect at 60°C, which presumably leads to irreversible destabilization of LPL.

**Fig. 7:**
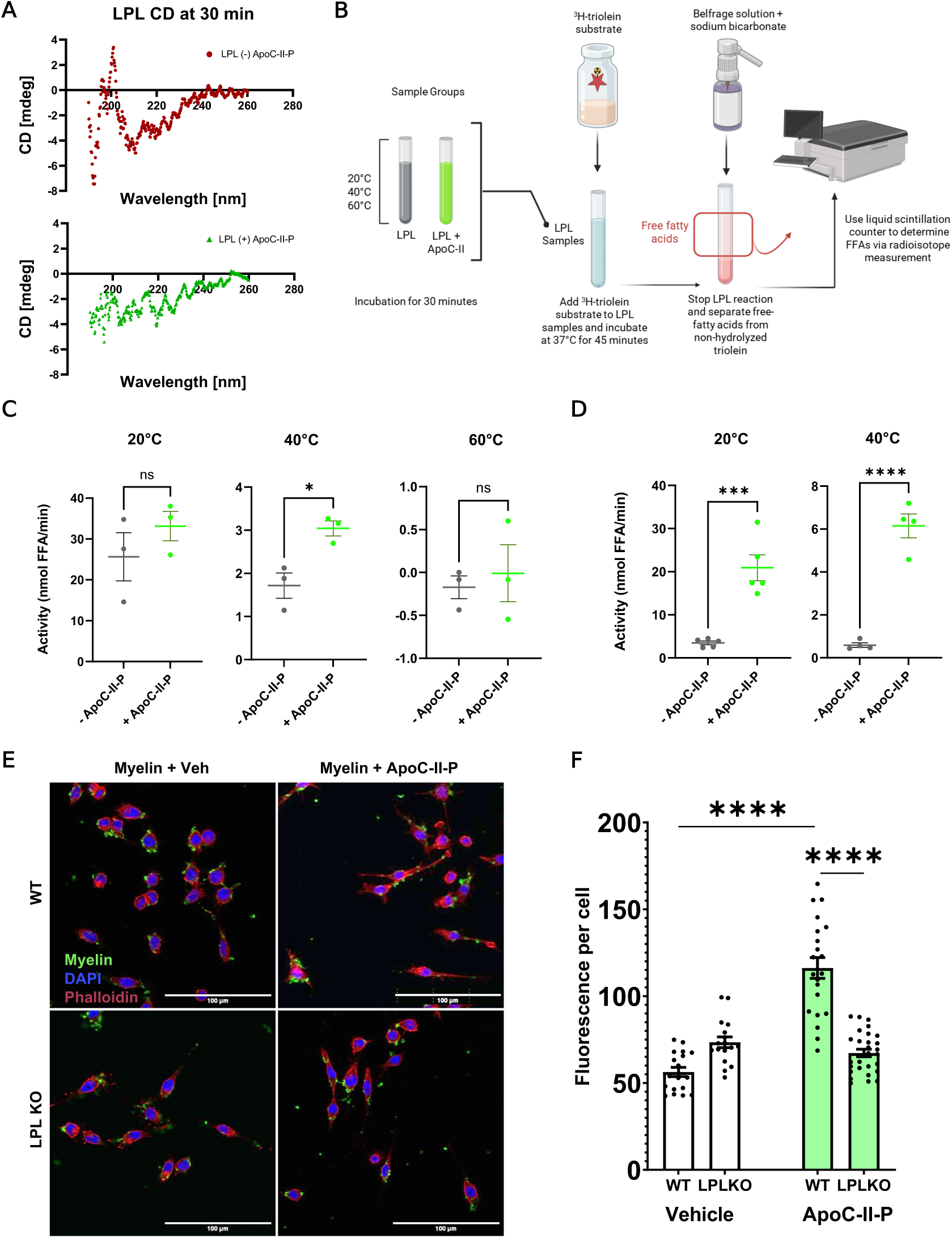
ApoC-II-P mitigates heat-induced inactivation of LPL and promotes LPL-mediated phagocytosis. (A) Circular dichroism (CD) spectra of bLPL (±) ApoC-II-P collected at 30 minutes, at 20°C (n=1). (B) Schematic of the LPL activity assay. LPL samples, with and without ApoC-II-P, are incubated at designated temperatures. Samples are then exposed to an LPL substrate containing radioactive (H3) triolein +/- ApoC-II-P. Following triolein/triglyceride hydrolysis, the Belfrage method is used to separate free fatty acids (FFAs) from non-hydrolyzed triglycerides. Radiolabeled FFAs, a direct readout of LPL-mediated hydrolysis, is then quantified by a liquid scintillation counter. (C) LPL is preincubated with 5 μM (+ApoC-II-P) for 30 minutes at 20°C, 40°C, and 60°C; 5 μM ApoC-II-P is also present in the LPL substrate. LPL activity is significantly lower at 40°C compared to 20°C, but ApoC-II-P partially mitigates heat- and time-associated loss of activity at 40°C. At 60°C, hydrolytic activity is lost in both conditions (n=3). Student’s t-test was used for significance (* = p<0.05). (D) LPL is preincubated with 5 μM ApoC-II-P (+ApoC-II-P) for 30 mins at 20°C (n=5) and 40°C (n=4) but is not added to the substrate. LPL activity is significantly higher when samples are incubated with ApoC-II-P at both temperatures. Student’s t-test was used for significance (*** = p<0.001, **** = p<0.0001). (E) WT and LPL KO BV2 cells were exposed to 10 mg/ml CFSE-labeled myelin (green) +/- 5 μM ApoC-II-P for 1 hour at 37 °C. Nuclei were labeled with DAPI (blue), and F-actin was labeled with Phalloidin (magenta). Myelin phagocytosis was increased in WT cells treated with ApoC-II-P. In contrast, no increase in phagocytosis was observed in LPL KO cells. (F) ImageJ quantification of CFSE-labeled myelin per cell (n=18,17,21,28; left to right, resp.). One-way ANOVA with a post-hoc Tukey’s test was used for significance (**** = p<0.0001).

To test if adding ApoC-II-P to the substrate partially rescued LPL activity following heat challenge, we conducted a second multi-temperature experiment where ApoC-II-P was not added to the substrate but instead was only incorporated during the 30-minute temperature-incubation period. Only the 20°C and 40°C conditions were tested, since all LPL activity was lost 60°C in the previous test. As before, hydrolytic activity was greatest following pre-incubation at 20°C (Fig. 7D). However, the samples pre-incubated at 20°C and 40°C in the presence of ApoC-II-P showed higher LPL activity (Fig. 7D), in contrast to our previous observations with APOC-II-P in the substrate (Fig. 7C), further supporting the idea that ApoC-II-P could rescue partial destabilization following room temperature incubation periods. Overall, these findings demonstrate that the ApoC-II-P can mitigate the heat- and time-associated loss of LPL’s hydrolytic activity, consistent with the maintenance of lid flexibility and N and C-terminal domain tethering observed in the MD simulations.

Next, we asked whether ApoC-II-P could modulate the function of cells that abundantly express LPL. Since BV2 microglial cells abundantly express LPL^4,5,42^, we used CFSE-labeled myelin debris to determine wither ApoC-II-P could alter the cells ability to phagocytose myelin debris. Moreover, to determine whether an ApoC-II-P-mediated change in phagocytosis was LPL-dependent, phagocytosis assays were performed in both wildtype (WT) and LPL knockout (LPL KO) cells^5^. Cells were supplemented with 5 μM ApoC-II-P for 1 hour before exposure to myelin debris. Image analysis revealed an increase in internalization of myelin debris in WT cells pre-treated with ApoC-II-P group (Fig. 7E-F). In contrast, LPL KO cells did not show an increase in myelin uptake when supplemented with ApoC-II-P (Fig. 7E-F). Notably, LPL KO cells exhibited a more ‘rod-like’ morphology, indicative of inflammatory polarization. These findings support the notion that ApoC-II-P enhances LPL-mediated phagocytosis of extracellular debris, and suggest that the ApoC-II-P peptide also improves the non-hydrolytic functions of LPL. This is also consistent with our MD analysis, which shows that ApoC-II-P prevents CTD flipping, which would alter the accessibility and orientation of the lipoprotein receptor binding domain, impairing the ability of LPL to bridge lipid substrates and cell surface receptors through its non-canonical functions^14,15,36,37^.

## Discussion

LPL plays key roles in lipid metabolism, including triglyceride hydrolysis, phospholipid hydrolysis, and ligand/receptor bridging. ApoC-II is a well-known activator of LPL and critical for its activity, but the exact molecular mechanisms by which these two proteins interact have not been fully elucidated. Herein, we described the use of MD simulations to model LPL over long timescales, both with and without an ApoC-II mimetic peptide, in conjunction with lipase activity experiments. Our simulations showed that LPL’s lid can access open, intermediate, and closed states. Furthermore, we observed a novel mechanism by which LPL destabilizes via CTD flipping, which is when the LBS and LRP-BS flip upside down. These structural motifs altered multiple LPL functional domains by shifting the N- and CTDs away from one another, reducing H-bond capabilities within catalytic triad atoms, and decreasing accessibility of the LRP-BS. Importantly, we discovered that ApoC-II-P did not anchor LPL’s lid but rather promoted lipid flexibility, consistent with the association between lid flexibility and LPL activity. ApoC-II-P did, however, promote stability via an alternative mechanism: bridging LPL’s N- and CTDs, which was more frequent when ApoC-II-P initially contacted LPL with an open lid. These findings were supported by biochemical assays, which showed that ApoC-II-P increased the thermostability of LPL. Finally, cellular assays revealed that ApoC-II-P can also promote the non-hydrolytic functions of LPL, such as phagocytosis.

Our simulations are the most extensive all-atom MD sampling of LPL to date, providing us with novel insights into LPL conformations and behavior over robust computational timescale and replicates. Further, due to the nature of our simulations, we were able to study LPL’s lid dynamics for the first time computationally. This allowed us to categorize the open, intermediate, and closed lid states, which had not been previously recognized due to the flexible nature of this region preventing its crystallization in all but one published LPL structures. A consistent observation throughout our study is that the intermediate lid state is the most frequently sampled among the LPL simulations (64.68% in No ApoC-II-P; 75.40% with ApoC-II-P). This may be in part due to the lid’s sensitivity to nearby ligands and its ability to discriminate them. The lid may change its position in the presence of specific lipids or lipoprotein components, activators like ApoC-II, or inhibitors like angiopoietin-like 4 (Angptl4)^30^. Thus, we naturally anticipate that this intermediate state is most frequently sampled, as the lid constantly transitions from open to closed, back to open. This self-regulation may imply how LPL transitions from active to inactive states, as the lid controls pore accessibility. Moreover, the presence of intermediate and open lid states likely associates to enzymatically active conformations.

The length of our simulations also allowed us to observe a never-before-seen behavior in LPL, which we described as CTD flipping. Seeing as these flips occurred after extensive sampling time, they likely reflect a conformational mechanism in which LPL inactivates due to its unstable nature over time and lack of CTD-specific binding partners (e.g., lipoprotein receptors). Our theory that flipping inactivated LPL was illustrated through several changes in functional domains. Notably, one of these was in the mediation of pore accessibility, reflected in catalytic triad residues. We noted that only the serine-histidine distance changed with CTD flipping. While the aspartate-histidine bond is generally recognized as the necessary H-bond in catalytic triads, the serine-histidine bond further supports histidine as a strong nucleophile and creates the charge-relay system. In terms of LPL’s hydrolytic activity, perhaps the tuning of this functional domain is largely dependent on this serine-histidine bond due to its reactivity across lid orientation and flipping as an on/off switch. Further, seeing as flipping consistently decreased the accessibility of the LRP-BS, we believe that this phenomenon prevents destabilized, flipped LPL from binding with receptors at the cell surface. Perhaps flipping as a whole is a regulatory mechanism for LPL, where it prevents unstable LPL from binding to lipids and receptors to ensure that active, stable LPL can properly perform hydrolysis and bridging with the limited number of necessary ligands and receptors. This aligns with our observation that the LBS maintained α-helicity with initial open lid configurations that proved to be an optimal interface for oleic acid binding, whereas closed and intermediate lids had reduced structure in this region. The promotion of increased lipid binding and hydrolysis suggests a unique structural relationship between the lid and LBS, and perhaps the greater CTD.

These observations hold true in the presence of ApoC-II-P. Binding of ApoC-II-P did not alter these structure-function trends across LPL, suggesting that the peptide promotes LPL activity by an alternative mechanism. Indeed, we found that when ApoC-II-P makes its initial contact with an open-lid LPL, it is able to bind between LPL’s NTD and CTD. This prevents CTD flipping and thus promotes an active form of LPL. Further, we noted that in some cases, the bridging enabled the formation of a continuous interface from the LBS to the lipid pore. Perhaps ApoC-II-P can achieve a greater functionality for transporting lipids across LPL for hydrolysis, as its charged residues strongly bind to LPL and thus leave its hydrophobic residues free for lipid binding. The nature of this interface also maintained LPL lid flexibility, which we believe is critical for normal function. While ApoC-II-P does interact with the lid, too strong of binding may anchor the lid into suboptimal positions that are not dynamic enough for lipid hydrolysis and bridging to occur, suggesting a noncanonical role for our specific ApoC-II peptide that is unique from full-length ApoC-II.

The ability of ApoC-II-P to bridge LPL’s N- and C-termini while maintaining lid flexibility may, in part, stem from its lack of stable secondary structure. Our simulations and circular dichroism spectra confirmed that ApoC-II-P does not adopt a helical conformation, yet it still enhances LPL activity in both hydrolytic and myelin-uptake functions. This is particularly intriguing given prior experimental and simulation studies demonstrating that ApoC-II peptides with stable helices promote LPL activation^43–45^. The mechanism by which our non-helical ApoC-II-P retains this activating capacity remains unclear but may relate to its sequence composition. Specifically, ApoC-II-P includes the conserved residues S78–A80 within the LPL-activating region, whereas other peptides^43–45^ feature substitutions at these sites. However, the helical structure of ApoC-II peptides may be necessary to mediate LPL-lipid binding and prevent self-aggregation. Further studies dissecting the chemistry and structural context of this ApoC-II LPL-binding domain could clarify its interactions with LPL and guide the design of shorter, optimized peptides with improved activity for therapeutic use.

MD simulations do not come without limitations. Here, protein docking is limited to nonflexible interfaces and may not represent realistic binding interfaces, but we believe that our long timescale simulations, in triplicate, overcome this barrier. Further, our experimental assays use bLPL while the simulations use human LPL (hLPL). However, we have previously shown that this ApoC-II-P can profoundly activate hLPL found in post-heparin plasma^34^, and given the structural similarity and availability of the bLPL, we believe these assays are representative and physiologically relevant. We also recognize that our simulations do not include other components that may regulate LPL function. For example, LPL is N-glycosylated at two sites (marked in Fig. 1). Due to the computational complications surrounding simulating glycans and that these sites were not near the lipid pore, lid, and CTD binding sites, we chose to simulate LPL without them. LPL also binds Ca^2+^ and other proteins, like itself, GPIHBPI, lipoprotein receptors, and heparan sulfate proteoglycans, that mediate stability in its CTD^16–18,21,46^. Considering that these likely underlie dimerization, and our simulations involve monomeric LPL, we chose not to include them to accurately model monomeric LPL that may or may not be soluble, but this may form the basis of future studies. Further, LPL and ApoC-II-P structure may change in the presence of lipids. Ultimately, simulations must include more expansive environments that include lipids, membranes and their embedded receptors, and other soluble proteins if we are to fully understand LPL activity. However, since our goal here was to understand how our ApoC-II mimetic peptide promotes LPL activity, we believe that our simulations accurately describe these mechanisms that cannot be attributed to other protein interactions.

LPL mediation shows promise as a potential therapeutic that would ultimately drive shifts in lipid metabolism across neurodegenerative and cardiovascular diseases. However, in order to develop effective therapeutics, we must have a detailed understanding of how LPL modulates multiple roles through self-regulation and its interactions with other proteins, lipids, and cofactors. To this end, our work has elucidated multiple mechanisms in which LPL is able to alter its functional domains to promote or inhibit hydrolytic activity, lipid binding, and receptor binding. Furthermore, it has deepened our understanding in how cofactors like ApoC-II, and their derived peptides, influence LPL stability and structure across time- and heat-associated losses. The dynamics of LPL have yet to remain fully uncovered, but this work continues to help piece together how LPL can be modulated on a molecular level with therapeutic peptides to rescue lipid metabolism changes.

## Methods

### Structural Model Preparation

The AlphaFold^47^ structures of human LPL and ApoC-II were retrieved from the AlphaFold Protein Structure Database with the codes AF-P06858-F1^48^ and AF-P02655-F1^49^, respectively. The LPL and ApoC-II AlphaFold structures comprise the entire protein domains (residues M1-G475, numbered 1-475; and M1-E101, respectively). However, these structures contain the signal peptide domain, which is cleaved prior to protein-protein interactions. Thus, these regions (M1-A27 and M1-G22) was removed from LPL and ApoC-II, respectively. For the ApoC-II peptide (ApoC-II-P), the full protein structure was clipped to only contain the peptide portion (S78-E100; numbered 23-101), defined previously by Oldham et al^34^.

The AlphaFold LPL structure was used instead of existing crystal structures due to the uncertainty involving the lid domain. There are currently four published human monomeric LPL structures and two bovine LPL structures, one dimeric^21^ and the other helical^18^. The four human LPL crystal structures are all bound to glycosylphosphatidylinositol-anchored high density lipoprotein-binding protein 1 (GPIHBP1), as well as a variety of unique ligands and/or branched monosaccharides. Of these four, three^16,17^ have unresolved lid and lipid binding domains. One model has these regions resolved, but there are small ligands bound to the lipid pore and lid domain^17^. Thus, to not bias any of the potential lid dynamics, we chose to simulate the AlphaFold structure.

### Pre-Docking MD Simulations

All molecular dynamics simulations were carried out using GROMACS 2023 software^50^. Proteins and ions were represented with the CHARMM36 force field^51^, and water molecules were modeled with the TIP3P-modified forcefield^52^. Each system first underwent energy minimization in vacuum, followed by a second minimization step after solvation and ion neutralization to ensure structural relaxation. We then equilibrated the systems under NVT conditions for 1 ns at 310 K using a modified Berendsen thermostat^53^. This was followed by a 1 ns NPT equilibration phase at 310 K and 1 bar, employing the same thermostat together with the Berendsen barostat^53^. Production simulations of LPL and ApoC-II-P were run independently in triplicate for 1 µs each, yielding a total of 6 µs of sampling prior to docking. All production simulation trajectories were generated using the Berendsen thermostat^53^ and the Parrinello–Rahman barostat^54^ maintained at 310 K and 1 bar, respectively. A 2-fs timestep was used, with full periodic boundary conditions and a 1.4-nm neighbor list cutoff. Bond constraints involving hydrogen atoms were enforced with the LINCS algorithm^55^. Long-range electrostatics were computed with the particle mesh Ewald (PME) method^56^ using a 1.2-nm real-space cutoff, and van der Waals interactions were cutoff at 1.2 nm.

### Selection of Structures for Docking

To evaluate structural stability and convergence during the production simulations, the root-mean-square deviation (RMSD) of each protein’s Cα atoms was calculated over time, using the corresponding energy-minimized structure as the reference. We also computed the root-mean-square fluctuation (RMSF) of the Cα atoms for each protein, averaging these values over the last 100 ns of each simulation. As with the RMSD analysis, the energy-minimized structure served as the reference for all RMSF calculations.

For LPL, we sought to create various conformations dependent on the dynamics of the lid region. Lid distance was defined as the distance between the Cα atoms of P187-G256. These non α-helical residues were chosen to evaluate how flexible and “open” the lid was in reference to its adjacent region near the lipid pore. Using GROMACS clustering tools^50,57^, clustering was performed−based on lid distance−to explore the dynamic conformations of LPL that were dependent on lid orientation. Mean lid distance and the standard deviation were computed across the entire simulation period for all LPL replicates in GROMACS. These values were used to cluster LPL conformations into three groups: open lid (distances more than one standard deviation above the mean), intermediate lid (distances within the mean ± one standard deviation), and closed lid (distances more than one standard deviation below the mean). Representative structures from each cluster (LPL-O, LPL-I, and LPL-C; open, intermediate, closed) were selected for docking.

For ApoC-II-P, the same clustering algorithm^50,57^ was utilized to identify the most representative structure across the three replicates, using the last 100 ns of each simulation. This structure was selected for docking analyses with LPL. The secondary structure motifs for both the pre-MD and post-MD, representative ApoC-II-P structures were calculated using the dihedral angles analysis module in the MDAnalysis Python library^58^.

### Metric and Cluster Definitions

All simulations were performed in triplicate, denoted by the R1-3 throughout the work described here. Simulations of LPL, conducted with and without ApoC-II-P, resulted in a number of LPL-lid-dependent structures and LPL-ApoC-II-P clusters. Herein, we describe the simulation sets and replicate names, and the specific structures used in each below in Table 1.

### Protein-Protein Docking

For rigid protein-protein docking, the online docking server ClusPro 2.0^59^ was employed. Three separate docking simulations were conducted for the following structures: LPL-O, LPL-I, and LPL-C, each paired with representative, average structure of ApoC-II-P. For each protein-peptide pair, 20 to 30 models were generated. Models were assessed for biological relevance in Visual Molecular Dynamics (VMD)^60^, determined by analyzing known LPL ApoC-II binding residues (aa77-87, aa114-126, aa212-244, aa245-263)^30^ within 5.0 Å of ApoC-II-P residues. These identified LPL residues were reported by Kumari et al. via hydrogen–deuterium exchange/mass spectrometry (HDX-MS) LPL-ApoC-II experiments^30^. ApoC-II residues known to activate LPL (Y85, I88, D91, Q92) within the LPL-activating region (S78-E101) via site-directed mutagenesis and lipase assays, reported by Shen et al., are contained within the peptide^29^. Thus, these LPL residues and all ApoC-II-P residues were used as a filter, such that the top models with the highest percentage of total LPL residues within 5.0 Å of ApoC-II-P residues were selected for post-docking MD simulations. In VMD, models were inspected to confirm there were not any docking artifacts. If any artifacts (spuriously interconnected loops) were present in models, those models were disqualified from consideration for simulations. Further, to ensure that steric clashes were not present in the docked models, a minimum distance calculation, between the residues of LPL and those of ApoC-II-P, was performed using the distances module in the MD Analysis Python library^58^. Models that were below a threshold of 2.5 Å were excluded from simulations.

### Protein-Lipid Docking

The online docking server SwissDock^61^ was used for protein-lipid docking, using the AutoDock Vina^62^ option which allowed for limited flexibility in the side chains of the lipid during docking. Six docking simulations were performed, using two LPL structures (LPL-O, LPL-C) and three lipids (oleic acid (OLA), phosphatidyl serine (SOPS), phosphatidyl choline (SOPC)). OLA structure was sourced from PubChem^63^, and SOPS and SOPC were created using the CHARMM-GUI server^64^. Lipid structures were converted to a Mol2 form using Open Babel^65^. Targeted docking on the LPL lipid-binding site (LBS) was utilized, with the search space being defined in a 30^3^ Å^3^ box surrounding the binding site. Swissdock^61^ generated 10 to 20 models for each protein-lipid pair. The top-scoring model was selected for visualization (in VMD^60^) and characterization. The distance module in the MD Analysis Python library^58^ was used for characterization of the model, including calculation of the number of residues within the LPL LBS that were within 5 Å of the lipid (binding interface); the number of hydrophobic residues (contacts) at this interface; and the number of potential H-bonds at the interface.

### Post-Docking MD Simulations

Each LPL-ApoC-II-P complex generated by ClusPro 2.0 was subjected to a 1-µs molecular dynamics simulation run, in triplicate, adding 9 µs of production sampling per pair (18 µs of simulations in total). The docked complexes were simulated using the same workflow and parameters outlined above for the pre-docking MD simulations.

### Simulation Trajectory Analysis

Model evaluation involved examining structural changes in LPL across the simulations. Standard GROMACS tools were used to compute the Cα RMSD and RMSF for both LPL and ApoC-II-P. In addition, the lid distance of LPL was determined throughout the trajectories using the same analysis approach described above.

The distance of the LPL lid to its lipid-binding site (LBS) were measured to quantify changes in LPL conformation due to C-terminal flipping upside down, with respect to the N-terminal. This distance was measured as the minimum distance from any lid residue (aa245-263) to that of LBS residues (aa412-422), using the distances module in the MDAnalysis Python library^58^. To identify pre- and post-flipping, a threshold of 33 Å was applied to the lid-LBS distances over time. Once LPL reached this distance, any simulation time at that point and on was categories as “post-flip,” and simulation trajectories were subsequently sorted.

Other distances to describe additional LPL domains were calculated using the methods described above. LPL lid to C-terminal domain (CTD) distance was quantified as the minimum between the lid (aa245-263) and the CTD (aa340-475). For lipid pore measurements, distances were calculated between the donor-acceptor pairs of the three bonds in the catalytic triad (S159:OG-H268:NE2, D183:OD1-H268:ND1, D183:OD2-H268:ND1).

To assess for changes in the availability of the LRP binding site (aa432-441), the solvent-accessible surface area (SASA) of this region was calculated using the GROMACS sasa algorithm^66^ over the entire simulation time. To further characterize the lipid pore^21^ identified by Gunn et al., the radius of gyration of residues lining the entire pore (aa74-103, 153-159, 265-282) was measured over simulation time, using the radius of gyration module in the MDAnalysis Python library^58^. To quantify potential bridging between the LPL NTD and CTD, via the lid, the number of H-bonds between the Lid-CTD were calculated using the GROMACS hbond algorithm (bond distance cutoff=0.35 nm; bond angle cutoff=30°) over the entire simulation time.

To test for significant differences across the groups, a one-way Kruskal-Wallis ANOVA was initially performed. To assess for differences across individual group comparisons, post-hoc Dunn’s test was used. The number of samples per group was calculated, and any groups that were sampled less than 1% of the total simulation time (per simulation set, e.g., LPL-Alone or LPL-ApoC-II-P simulations) was excluded from the plots.

Secondary structure assignments of LPL residues were computed using the Compute DSSP module from the MDTraj Python library^67^. Calculations were performed over the entire simulation time and plotted against time. Assignments were listed as α-helix (H), residue in isolated β-bridge (B), extended strand participating in β-ladder, 3/10-helix (G), π-helix (I), hydrogen-bonded turn (T), bend (S), and loop/irregular elements (C).

LPL-ApoC-II-P simulations were further grouped according to multiple constraints for the analyses described above. These simulations were first sorted into pre- and post-flip groups, based on the threshold described earlier. Secondly, the lid distance (described in **Selection of Structures for Docking**) of these two groups was calculated into open, intermediate, and closed distances. Finally, the “nearness” of ApoC-II-P to the top of the lipid pore and lid (aa 81-91, 159-161, 183-188, 210-214, 238-271) was determined with a minimum distance calculation, using the method described earlier. If ApoC-II-P was within 5 Å of this LPL selection, they were categorized as ApoC-II-P: Near, and if not, into ApoC-II-P: Far.

The LPL and ApoC-II-P residues within 5 Å of each other^68^, over the last 100 ns of each simulation, were calculated using the distances module in the MDAnalysis Python library^58^. Distances were plotted in contact frequency maps per replicate. These residues were compared to those identified from spectroscopy and lipase assays in existing literature^29,30^. LPL residues that interacted with ApoC-II-P for 50+% of the calculated sample time were listed. Contact frequency maps were used to cluster binding sites across the nine LPL-ApoC-II-P simulations.

Interactions between LPL and ApoC-II-P were evaluated by estimating the complexes’ interaction free energies using the molecular mechanics Poisson–Boltzmann surface area (MM/PBSA) approach. This analysis was carried out with the gmx_MMPBSA tool^69^ integrated into GROMACS and was applied to the final 100 ns of each trajectory (900–1000 ns). We additionally performed an energy decomposition analysis to identify residues contributing most strongly to binding and to quantify their interaction frequencies. For each LPL-ApoC-II-P complex, free energy values were computed at 0.5-ns intervals across the analysis window, yielding 201 energy estimates per simulation. Energies were plotted as boxplots with standard deviations, and one-way Kruskal-Wallis ANOVA with post-hoc Dunn’s test were performed for all groups. Decomposition profiles were plotted for the top 20 stabilizing residues in LPL and the top 5 residues in ApoC-II-P.

### Lipoprotein Lipase Assay

The radiometric quantification of LPL activity was performed using a recently optimized assay developed by our group^34^. In brief, we prepared an H^3^ triolein-based substrate, which typically contains an LPL activator, such as ApoC-II-P^34^. The substrate was incubated with a source of LPL, in this case bovine LPL (bLPL, Sigma, L2254-5KU), for 45 minutes at 37°C. To quantify LPL-mediated hydrolysis, the reaction was stopped with sodium bicarbonate, and non-hydrolyzed triolein was separated from free fatty acids (FFAs) using Belfrage extraction. Radiolabeled FFAs, a direct readout of LPL-mediated hydrolysis, were then quantified by liquid scintillation (Fig. 7B). Here, LPL activity assays were performed at three different temperature conditions: 20°C, 40°C, and 60°C. bLPL samples were prepared as a 1:100 dilution in Krebs Ringer Phosphate Buffer (KRP). Samples were prepared in triplicate for each condition (20°C, 40°C, and 60°C) with/without the addition of 5 µM ApoC-II-P and incubated for 30 minutes before the typical reaction time (45 minutes at 37°C). Once the 30-minute incubation period was completed, 100 μL of substrate (+/- 5 µM ApoC-II-P) was added to the samples for 45 minutes at 37°C. Changes in LPL hydrolysis, presented as mmol FFA generated per minute, were determined by performing a Student’s t-test between +/- ApoC-II-P for each temperature condition in GraphPad Prism.

### Myelin Phagocytosis Assay

Wild-type (WT) and LPL knock-out (LPL KO) murine microglia (BV-2) cells were generated as previously described^5^. Cells were seeded at 25,000 cells/well and allowed to attach at 37°C in 5% CO_2_ overnight. Before exposure, the media was aspirated and replaced with vehicle-containing (H_2_O) media or media containing 5 μM ApoC-II-P. After 1 hour, 10 mg/mL CFSE-labeled myelin debris was spiked into each well and incubated for 1 hour at 37 °C, 5% CO_2_. The cells were washed 3 times with sterile PBS +1% fatty acid-free bovine serum albumin (BSA) to remove non-engulfed myelin debris. Cells were fixed with room temperature 4% paraformaldehyde (in PBS) for 10 minutes. Cells were washed twice with PBS, and F-actin was stained using Phalloidin (Phalloidin 594, Thermo Fisher) for 1 hour at room temperature. Cells were rinsed twice with PBS, and the nuclei were stained with DAPI. Cells were imaged using an Olympus FV1000 laser scanning confocal. Images were analyzed using ImageJ. Specifically, cell masks were generated by outlining cell perimeters using the Phalloidin stain and quantifying CFSE/GFP fluorescence intensity within each cell. As presented, fluorescence intensity is indicative of internalized myelin and, hence, myelin phagocytosis. Changes in myelin phagocytosis were determined by performing a one-way analysis of variance (ANOVA) and an appropriately conservative post-hoc test (e.g., Tukey) for multiple comparisons.

### Circular Dichroism to Detect Secondary Structures

Circular dichroism (CD) spectroscopy was performed to determine the secondary structure of peptides by measuring the differential absorption of left- and right-circularly polarized light. CD quantifies the presence of α-helices, β-sheets, and random coils to determine peptide makeup. CD spectra were collected for the human ApoC-II peptide (ApoC-II-P) at both 20°C and 60°C to evaluate temperature-induced structural changes. Additionally, CD was performed on bovine lipoprotein lipase (bLPL) in the presence or absence of human ApoC-II-P to assess the impact of the peptide on LPL secondary structure.

### Human ApoC-II-P

Human ApoC-II-P was prepared by diluting the 1 mM stock human ApoC-II-P in tris(2-carboxyethyl) phosphine hydrochloride (TCEP HCl; GoldBio, 51805-45-9) phosphate-buffered saline (1X) 0.0067M (PBS) for a 0.3mg/mL final concentration. Using a Jasco J-815 (Serial No. A029361168) CD Spectrometer’s Spectra Manager software, high sensitivity spectra were captured and processed. The baseline was set using 1 mM TCEP/PBS, and ApoC-II-P was loaded into a high-precision cell 1 mm quartz glass cuvette. CD measurements were taken at 20°C, followed by gradual heating over one hour to 60°C, at which point a second scan was performed. For each scan, data was collected every 2 sec/wavelength from 190 to 250 nm, using a bandwidth of 1.0 nm.

### Bovine LPL ± Human ApoC-II-P

Bovine lipoprotein lipase (bLPL; Millipore Sigma, L2254-5KU) was diluted 1:4 in 100 mM sodium phosphate buffer. CD spectra were recorded using the Jasco J-815 spectrometer and software as described above. The baseline was set to 100 mM sodium phosphate buffer, and the bLPL sample was loaded into a high-precision, 1 mm quartz glass cuvette. Spectra were collected at 20°C at 10-minute intervals over a 30-minute period to monitor time-dependent conformational changes. For each scan, data was collected every 2 sec/wavelength from 190 to 260 nm, with a bandwidth of 0.2 nm. A subsequent experiment was conducted using bLPL supplemented with the human stock 1 mM ApoC-II-P as a 1:5 ratio under identical conditions, with spectra collected every 10 minutes for 30 minutes. For individual spectra and comparisons, see Supplementary Fig. 15.

## Data availability

AlphaFold^47^ codes of the LPL and ApoC-II structures utilized in this work are: AF-P06858-F1^48^ and AF-P02655-F1^49^. SOPC and SOPS structures were sourced from CHARMM-GUI^64^ and OLA from PubChem^63^. The molecular dynamics coordinates, trajectories and structures used in this study are available upon reasonable request to the corresponding authors.

## Code availability

Molecular dynamics simulations in this were performed with GROMACS 2023^50^. The CHARMM36 force field^51^ was used for all simulations to model proteins and ions, with the corresponding TIP3P-modified forcefield^52^ for water. Lipid structure file manipulation was performed with Open Babel^65^. Docking was performed with ClusPro 2.0^59^ and Swissdock^61^, with AutoDock Vina^62^. Simulations were conducted with standard GROMACS tools. GROMACS modules^50^ [rms, rmsf, sasa, hbond] and Python libraries [MDanalysis^58^, MDTraj^67^] were used for protein structural analyses. The gmx_MMPBSA tool^69^ was used for binding free energy calculations. Structural figures were rendered using VMD^60^. Python, BioRender, and GraphPad Prism was used for graphical plotting. ImageJ (FIJI) was used for image analysis.

## Acknowledgements

This work utilized the Alpine high-performance computing resource and Blanca condo computing resource at the University of Colorado Boulder. Alpine is funded through a joint effort by the University of Colorado Boulder, the University of Colorado Anschutz, and Colorado State University. Blanca receives funding from its computing users and the University of Colorado Boulder. This work was supported by an ABNEXUS grant awarded to K.D.B. and K.G.S., a National Institutes of Health (NIH), National Institute on Aging (NIA) R01 awarded to K.D.B. (R01AG079217), and a NIH T32 awarded to E.E.L. (5 T32 DK 120520-5).

## Author information

### Authors and Affiliations

**Department of Chemical and Biological Engineering, University of Colorado Boulder, Boulder, CO, 80303, USA**

Emma E. Lietzke, Ziyue Dong & Kayla G. Sprenger

**Division of Endocrinology, Metabolism, and Diabetes, University of Colorado Anschutz Medical Campus, Aurora, CO, 80045, USA**

Emma E. Lietzke, Mary S. Rouse, Dean Oldham, Robert H. Eckel. & Kimberley D. Bruce

### Contributions

Conceptualization: E.E.L., K.G.S., K.D.B. Experimentation: M.S.R., D.O., K.D.B. Computational Modeling and MD Simulations: E.E.L. Simulation Analysis: E.E.L., Z.D. Writing−original draft: E.E.L., K.G.S., K.D.B. Writing−review & editing: E.E.L., M.S.R., D.O., Z.D., R.H.E., K.G.S.,

K.D.B. Revision: E.E.L., K.G.S., K.D.B. Supervision: K.G.S, K.D.B.

### Corresponding authors

Correspondence to Kayla G. Sprenger & Kimberley D. Bruce.

