## Supplementary Information for "Lipoprotein lipase requires a flexible lid and stable C-terminal, both maintained by ApoC-II peptide binding"

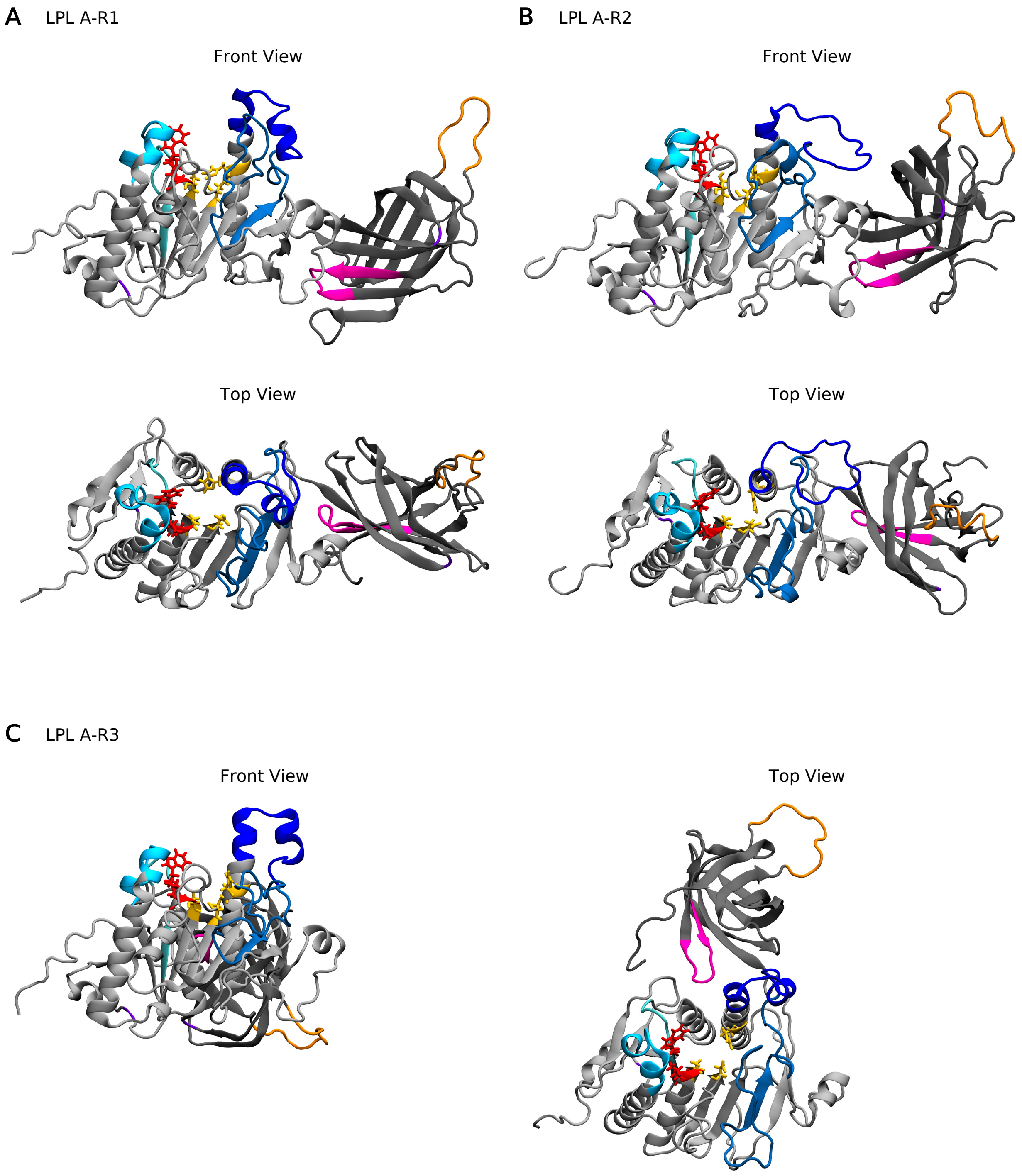


**Supplementary Figure 1. Final simulation poses of LPL.** LPL is represented as described in Fig. 1, with t=1000 ns. (A) A-R1 simulation. (B) A-R2 simulation. (C) A-R3 simulation.

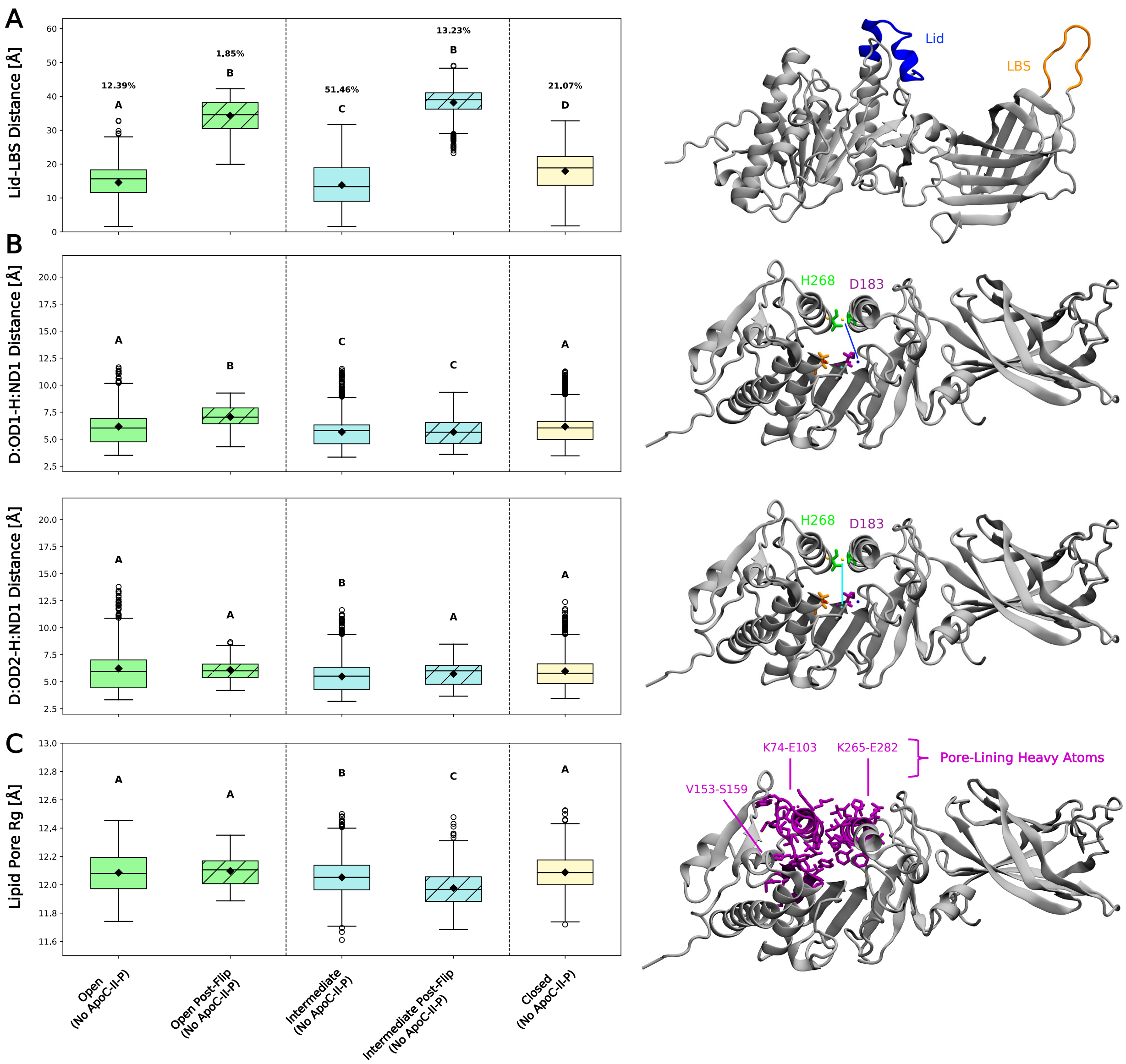


**Supplementary Figure 2. Lid orientation and CTD flipping impact on LPL functional domains.** Open lids are shown in green, intermediate in blue, and closed in yellow. Post-flip groups shown with hatching. Group occupation percentages, over total simulation time, are shown for each group. Compact-letter-display represents significantly different groups from a Kruskal-Wallis ANOVA with post-hoc Dunn's comparison tests (N=3 independent simulation runs, with 2,001 data points/run; see Supplementary Table 1 for n breakdown per group). (A) Left: Lid-LBS distance measured across groups. Right: Visualization of lid-LBS distance. (B) Left: Distance between the D183:OD1 and H268:ND1 atoms (top) and D183:OD2 and H268:ND1 atoms (bottom) across groups. Right: Visualization of the respective pore distances on the left. (C) Left: Radius of gyration (Rg) of residues lining the lipid pore across groups. Right: Visualization of the pore-lining residues, with only heavy atoms shown for visual clarity.

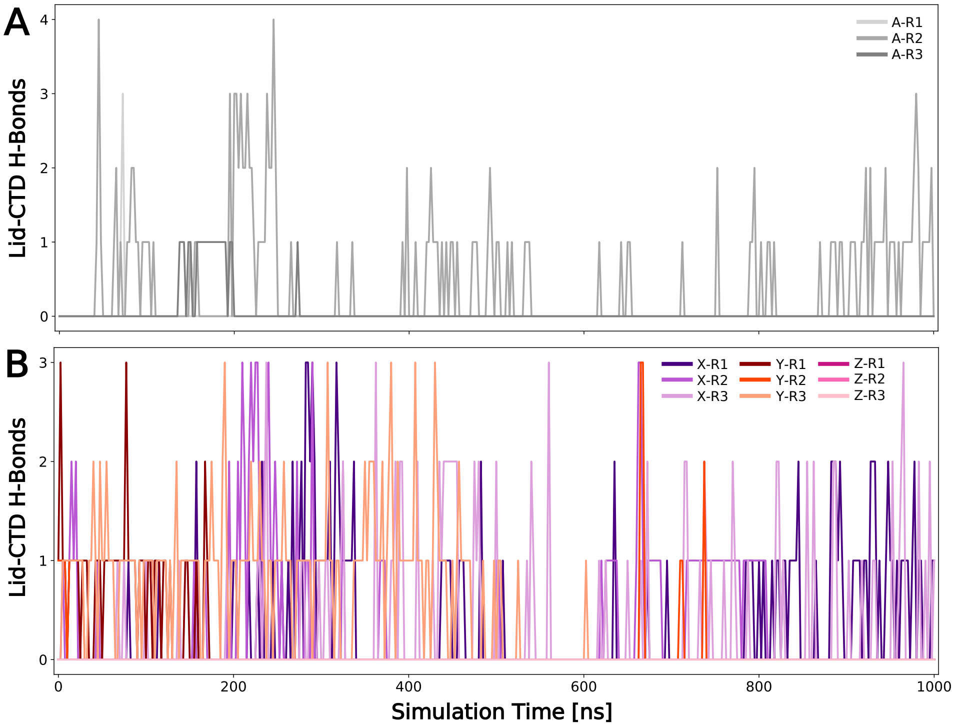


**Supplementary Figure 3. H-bonding between the lid and LBS, with the CTD of LPL.** Smoothed number of H-bonds between specified domains over simulation time. (A) Number of H-bonds between the lid and CTD for simulations without ApoC-II-P. Replicates are shown in gray. (B) Number of H-bonds between the lid and CTD for simulations with ApoC-II-P. Clustered colors (purple, orange, pink) represent various initial LPL + ApoC-II-P docking positions (X, Y, Z, resp.).

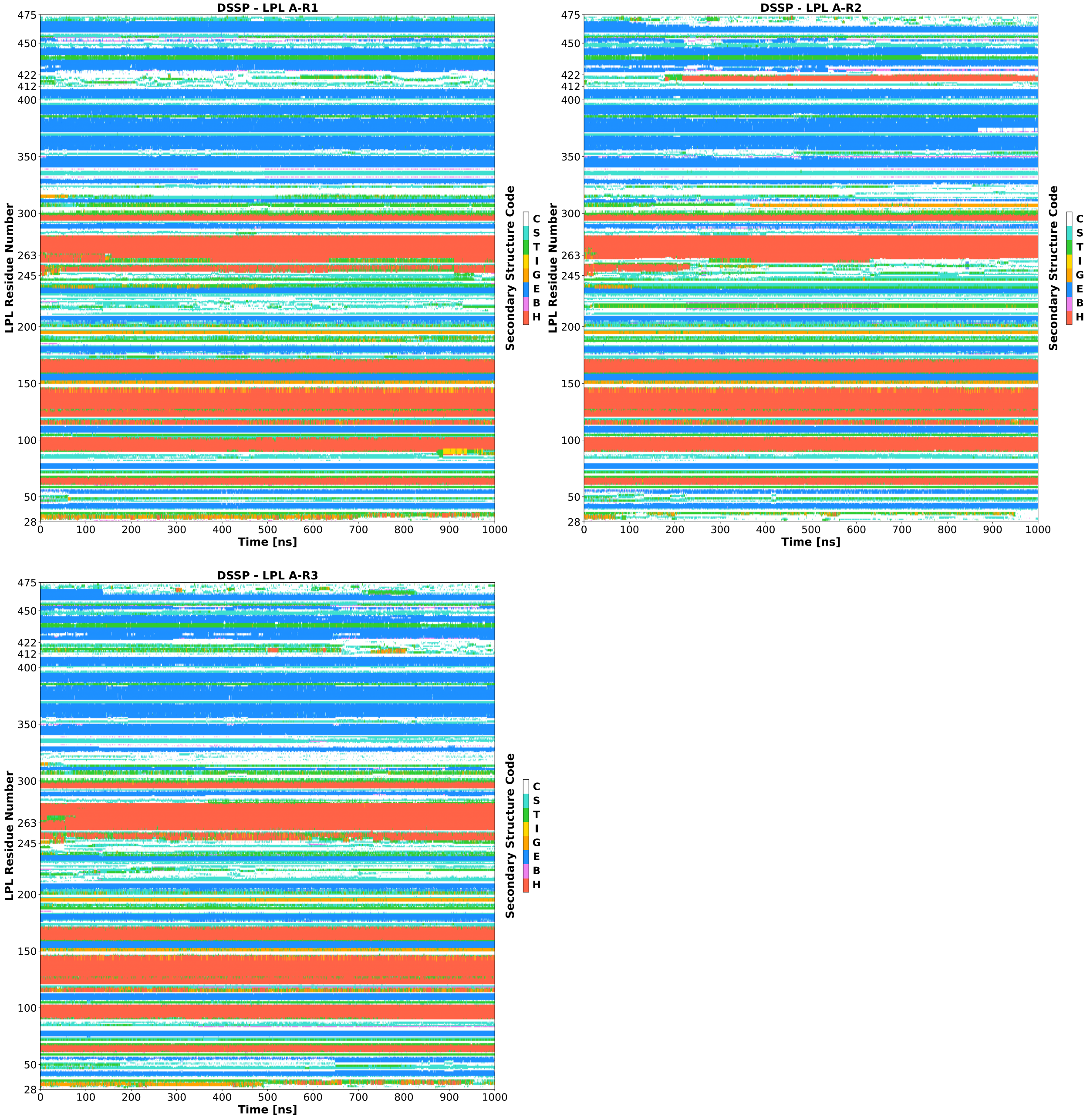


**Supplementary Figure 4. Dictionary of protein secondary structure (DSSP) assignments for A-R1, A-R2, and A-R3 per LPL residue number plotted against time.** Assignments codes are: α-helix (H), residue in isolated β-bridge (B), extended strand participating in β-ladder, 3/10-helix (G), π-helix (I), hydrogen-bonded turn (T), bend (S), and loop/irregular elements (C). Relevant residues for the LPL lid (aa245-263) and lipid binding site (aa412-422) are listed on the y-axis.

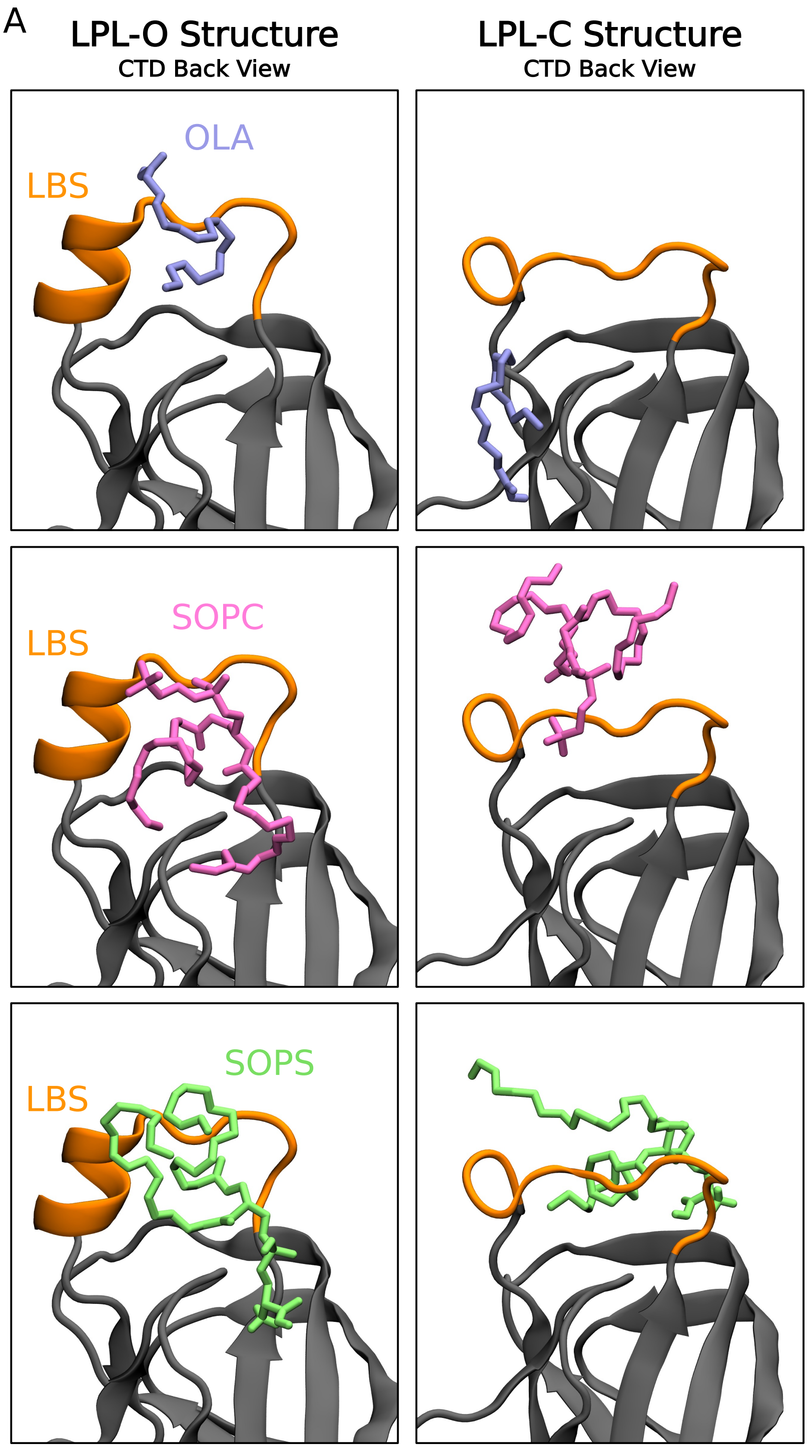


B

LPL LBS Contact %

| | LPL-O | LPL-C |
|---|---|---|
| OLA | 72.72 | 18.18 |
| SOPC | 63.63 | 72.72 |
| SOPS | 63.63 | 72.72 |

Hydrophobic Contact %

| | LPL-O | LPL-C |
|---|---|---|
| OLA | 11.59 | 0 |
| SOPC | 5.19 | 7.39 |
| SOPS | 5.90 | 6.88 |

Number of H-Bonds

| | LPL-O | LPL-C |
|---|---|---|
| OLA | 1 | 0 |
| SOPC | 0 | 2 |
| SOPS | 1 | 0 |

**Supplementary Figure 5. Lipid binding to variable LBS conformations of LPL.** (A) Visual representations of docking models of LPL with lipids. Models with LPL-O structures are seen on the left, and models with LPL-C structures are seen on the right. LPL's LBS is colored in orange. Top: Docking with oleic acid (OLA), seen in lavender. Middle: Docking with stearoyl-oleoyl-phosphatidylcholine (SOPC), seen in light pink. Bottom: Docking with stearoyl-oleoyl-phosphatidylserine (SOPS), seen in light green. (B) Contacts between the lipid and LPL LBS region were quantified for the six models shown in (A). Top: The percentage of residues in the LBS within 5 Å of the lipid. Middle: The percentage of hydrophobic contacts in the LBS with the lipid. Bottom: The number of potential H-bonds between LBS residues and the lipid.

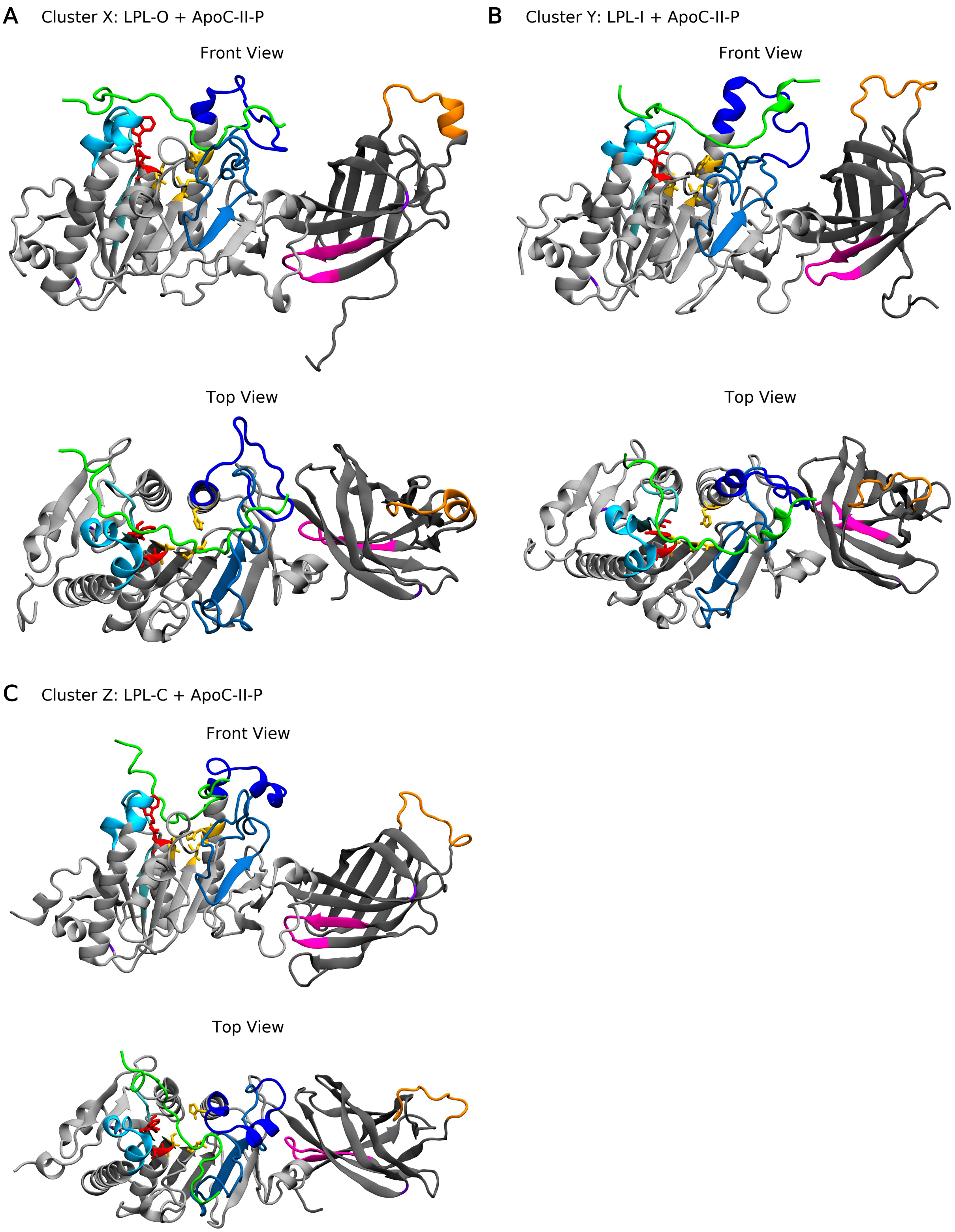


**Supplementary Figure 6. Initial docking poses selected for simulations.** LPL is represented as described in Fig. 1 with ApoC-II-P in green. (A) Starting point of the LPL-O and ApoC-II-P simulations, representing Cluster X simulations. (B) Starting point of the LPL-I and ApoC-II-P simulations, representing Cluster Y simulations. (C) Starting point of the LPL-C and ApoC-II-P simulations, representing Cluster Z simulations.

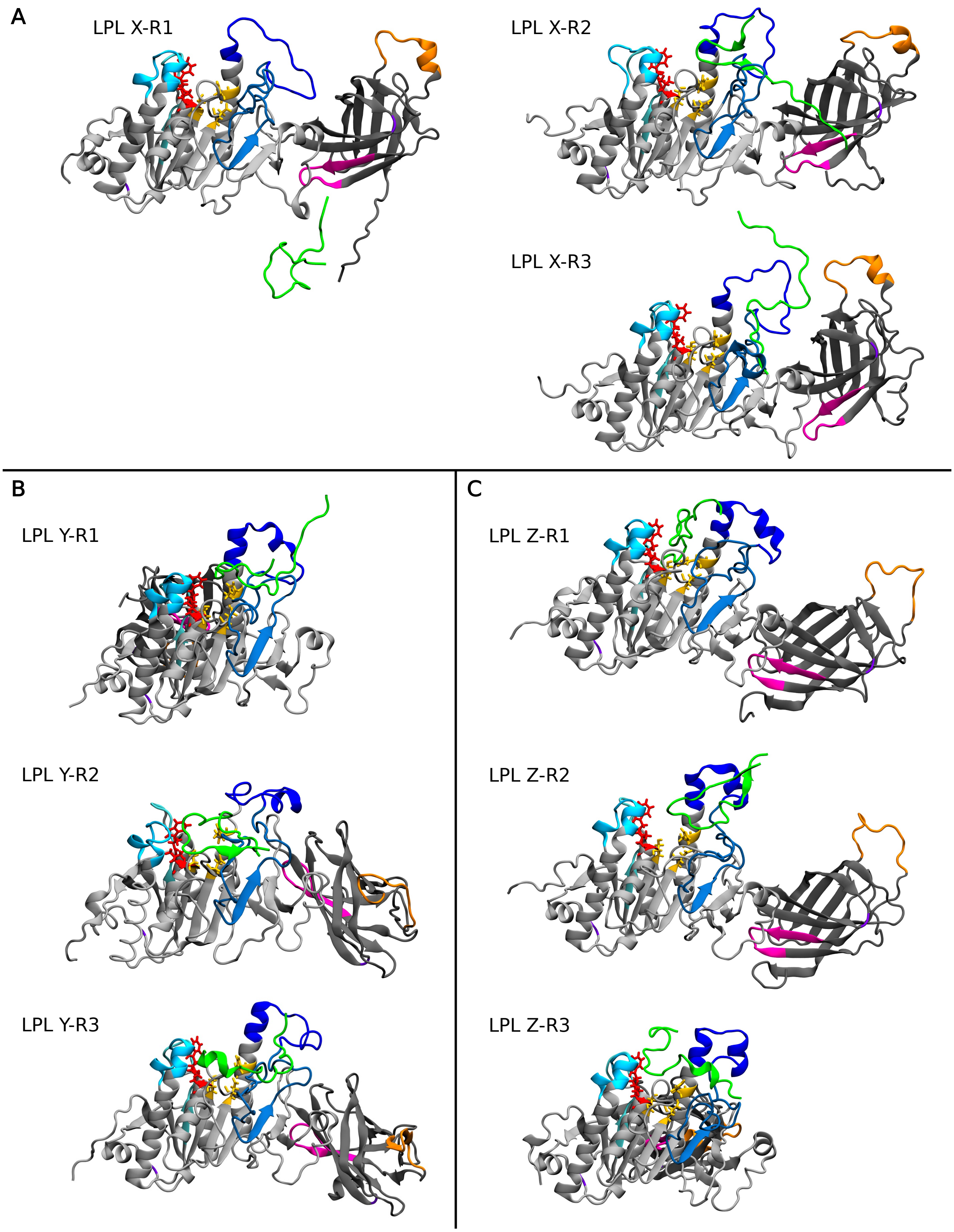


**Supplementary Figure 7. Final binding positions of LPL and ApoC-II-P.** LPL is represented as described in Fig. 1 with ApoC-II-P in green, t=1000 ns. (A) Cluster X simulations. (B) Cluster Y simulations. (C) Cluster Z simulations.

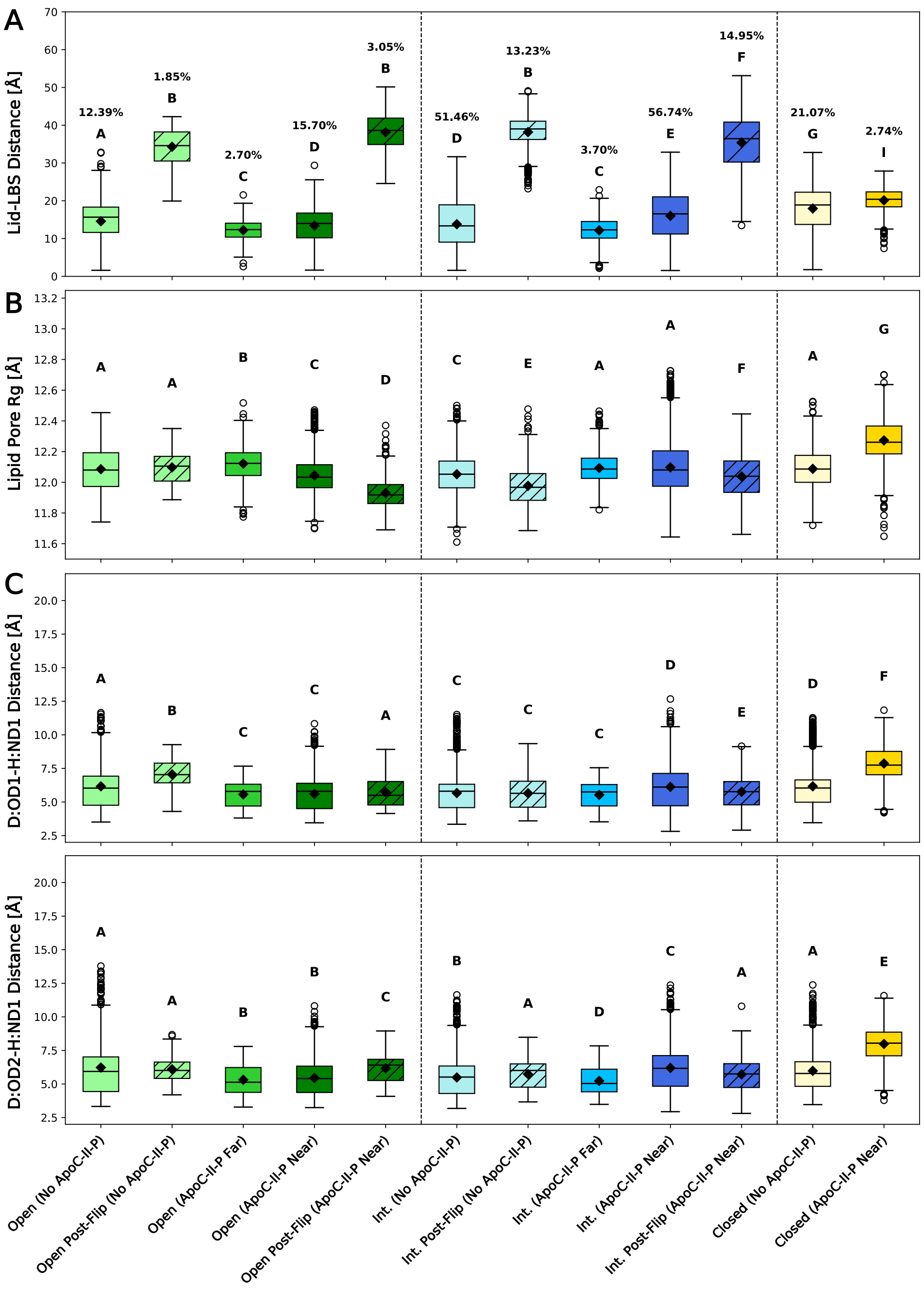


**Supplementary Figure 8. ApoC-II-P impact on functional domains of LPL.** Open lids are shown in green, intermediate in blue, and closed in yellow. Post-flip groups shown with hatching. Group occupation percentages, over total simulation time, are shown for each group (separate percentage totals were calculated for No ApoC-II-P and ApoC-II inclusive simulations). Groups represented less than 1% of the simulation time are not shown. Compact-letter-display represents significantly different groups from a Kruskal-Wallis ANOVA with post-hoc Dunn's comparison tests (N=12 independent simulation runs, with 2,001 data points/run; see Supplementary Table 1 for n breakdown per group). (A) Lid-LBS distance measured across groups. (B) Radius of gyration (Rg) of residues lining the lipid pore across groups. (C) Distance between the D183:OD1 and H268:ND1 atoms (top) and D183:OD2 and H268:ND1 atoms (bottom) across groups.

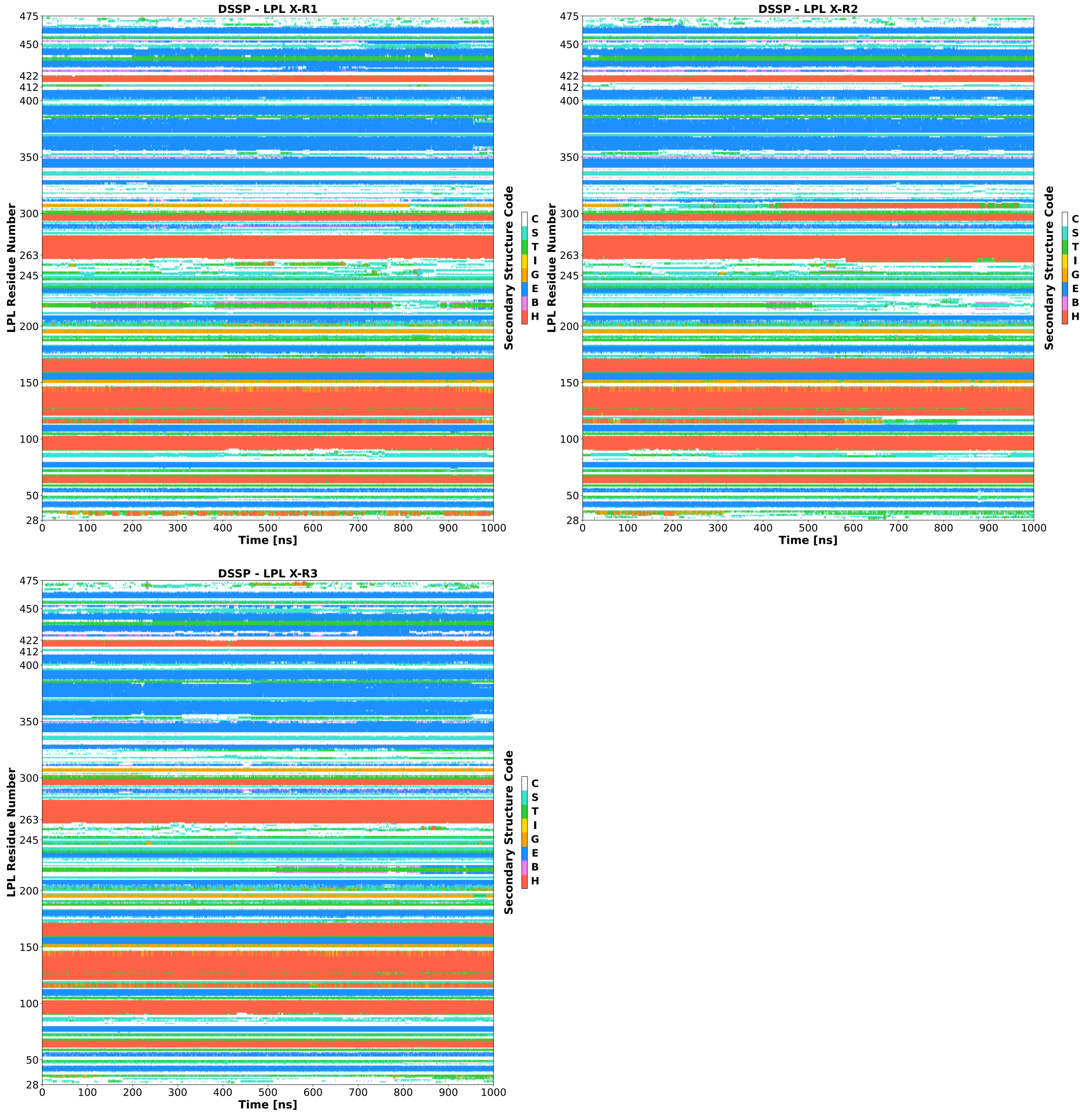


**Supplementary Figure 9. Dictionary of protein secondary structure (DSSP) assignments for X-R1, X-R2, and X-R3 per LPL residue number plotted against time.** Assignments codes are: α-helix (H), residue in isolated β-bridge (B), extended strand participating in β-ladder, 3/10-helix (G), π-helix (I), hydrogen-bonded turn (T), bend (S), and loop/irregular elements (C). Relevant residues for the LPL lid (aa245-263) and lipid binding site (aa412-422) are listed on the y-axis.

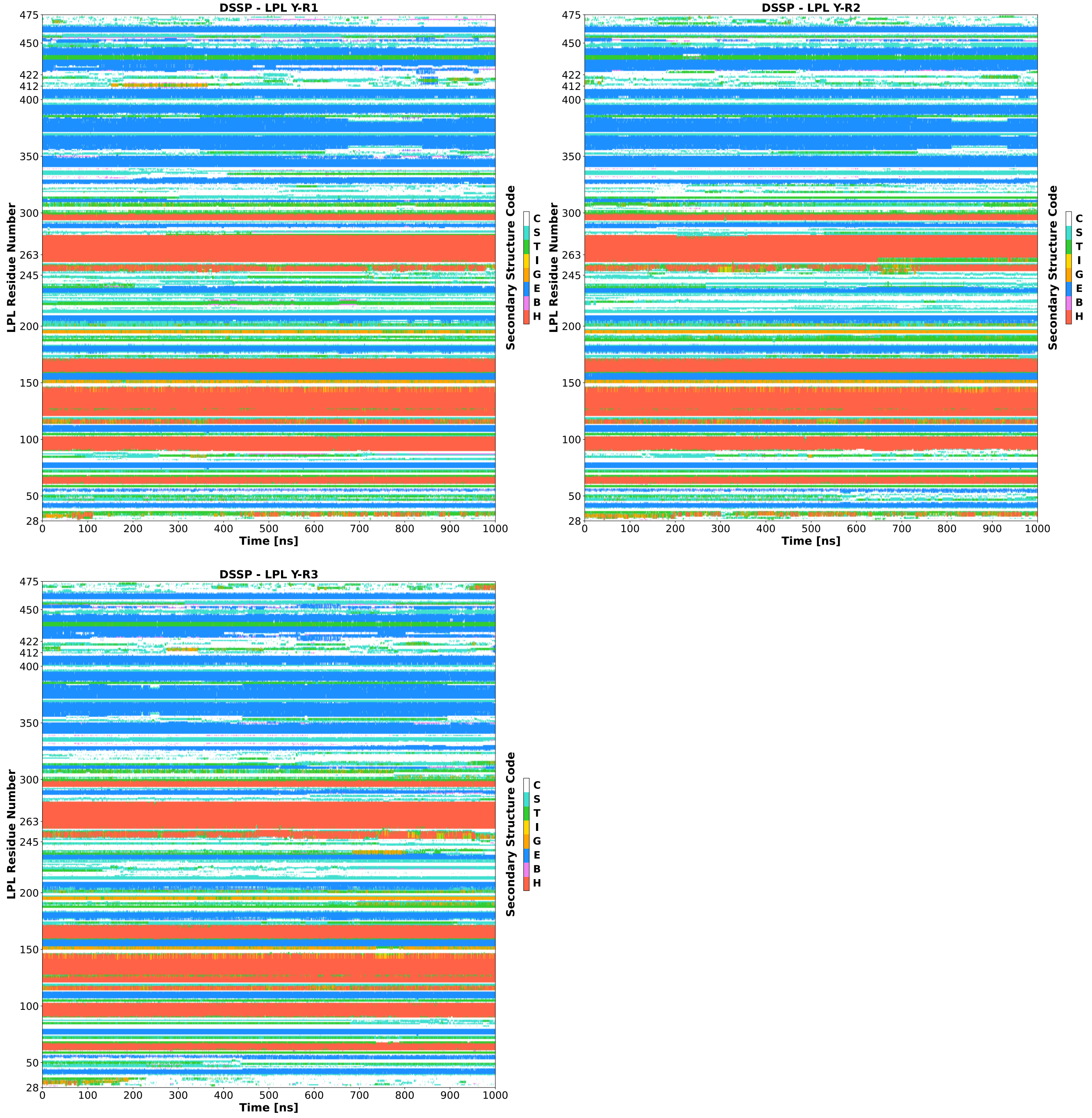


**Supplementary Figure 10. Dictionary of protein secondary structure (DSSP) assignments for Y-R1, Y-R2, and Y-R3 per LPL residue number plotted against time.** Assignments codes are: α-helix (H), residue in isolated β-bridge (B), extended strand participating in β-ladder, 3/10-helix (G), π-helix (I), hydrogen-bonded turn (T), bend (S), and loop/irregular elements (C). Relevant residues for the LPL lid (aa245-263) and lipid binding site (aa412-422) are listed on the y-axis.

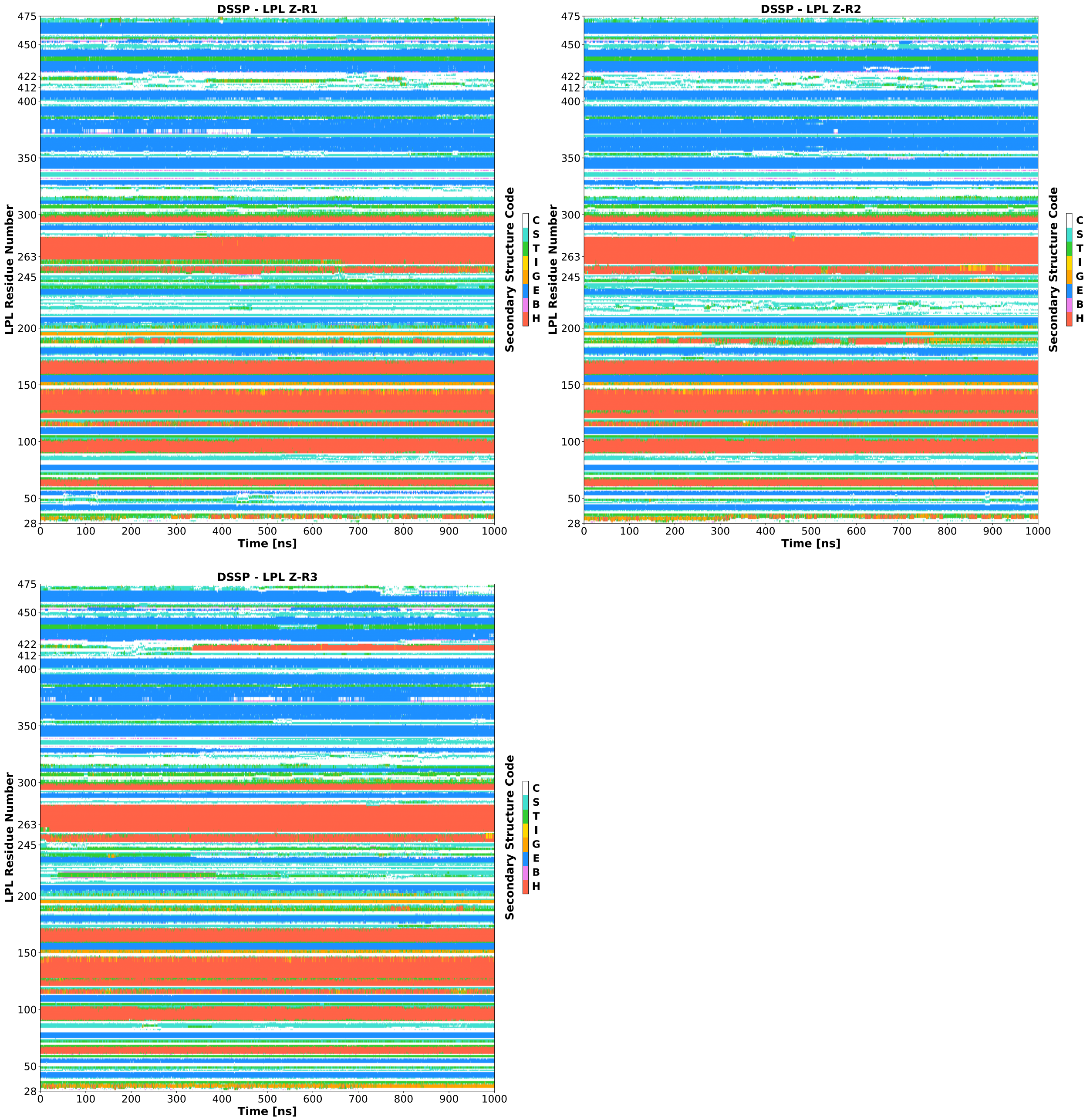


**Supplementary Figure 11. Dictionary of protein secondary structure (DSSP) assignments for Z-R1, Z-R2, and Z-R3 per LPL residue number plotted against time.** Assignments codes are: α-helix (H), residue in isolated β-bridge (B), extended strand participating in β-ladder, 3/10-helix (G), π-helix (I), hydrogen-bonded turn (T), bend (S), and loop/irregular elements (C). Relevant residues for the LPL lid (aa245-263) and lipid binding site (aa412-422) are listed on the y-axis.

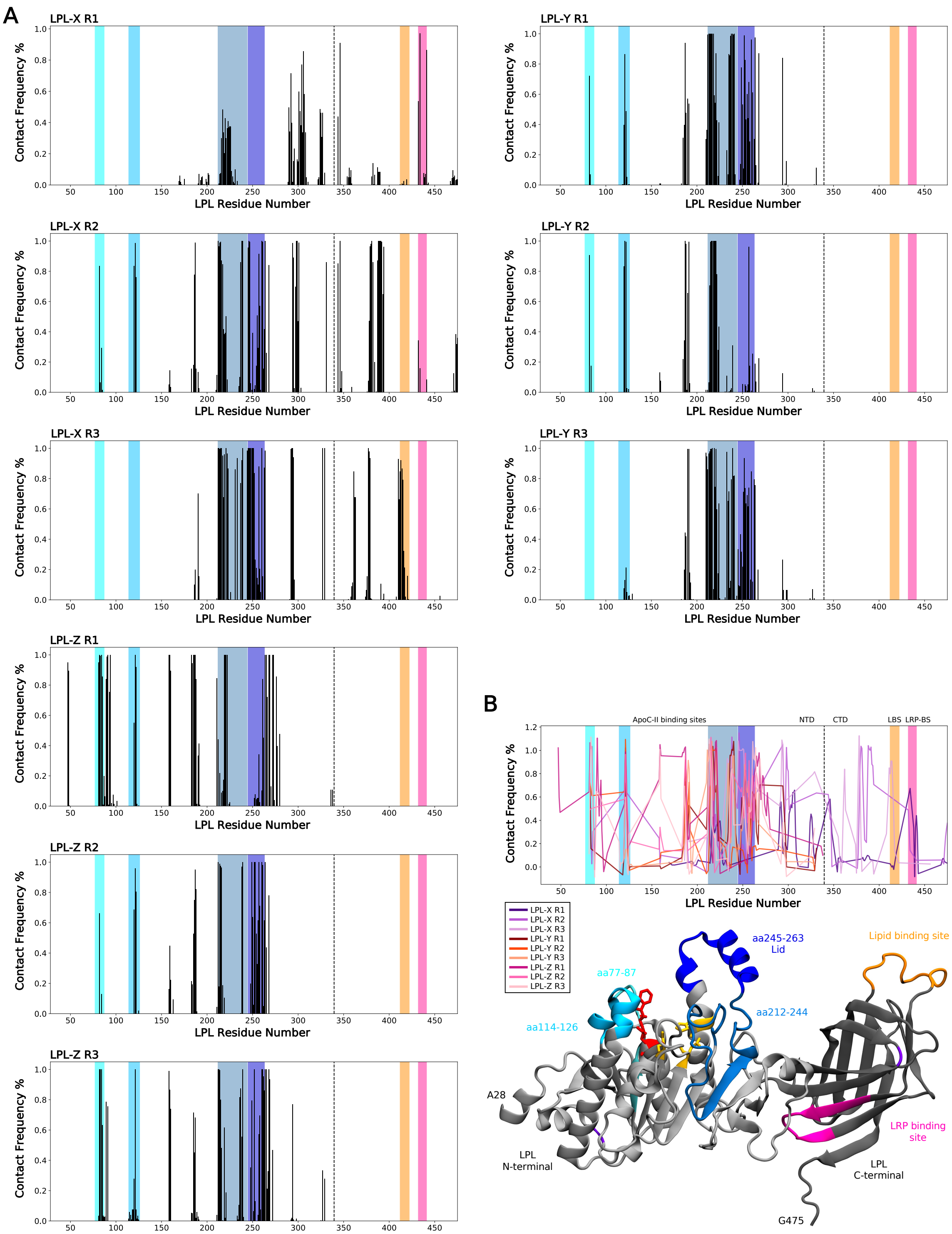


**Supplementary Figure 12. LPL residue contact frequencies with ApoC-II-P.** (A) The fraction percentage in which LPL residues were within 5 Å of ApoC-II-P over the final 100 ns of the trajectory was measured across simulations. Colored regions are identical to those in the RMSF graphs seen in Fig. 3C,5B, and also seen visually on LPL's structure and summary plot in (B). (B) Top: A smoothed, summary of contact frequencies across the nine simulations are overlayed. Bottom: LPL visualization, as seen in Fig. 1.

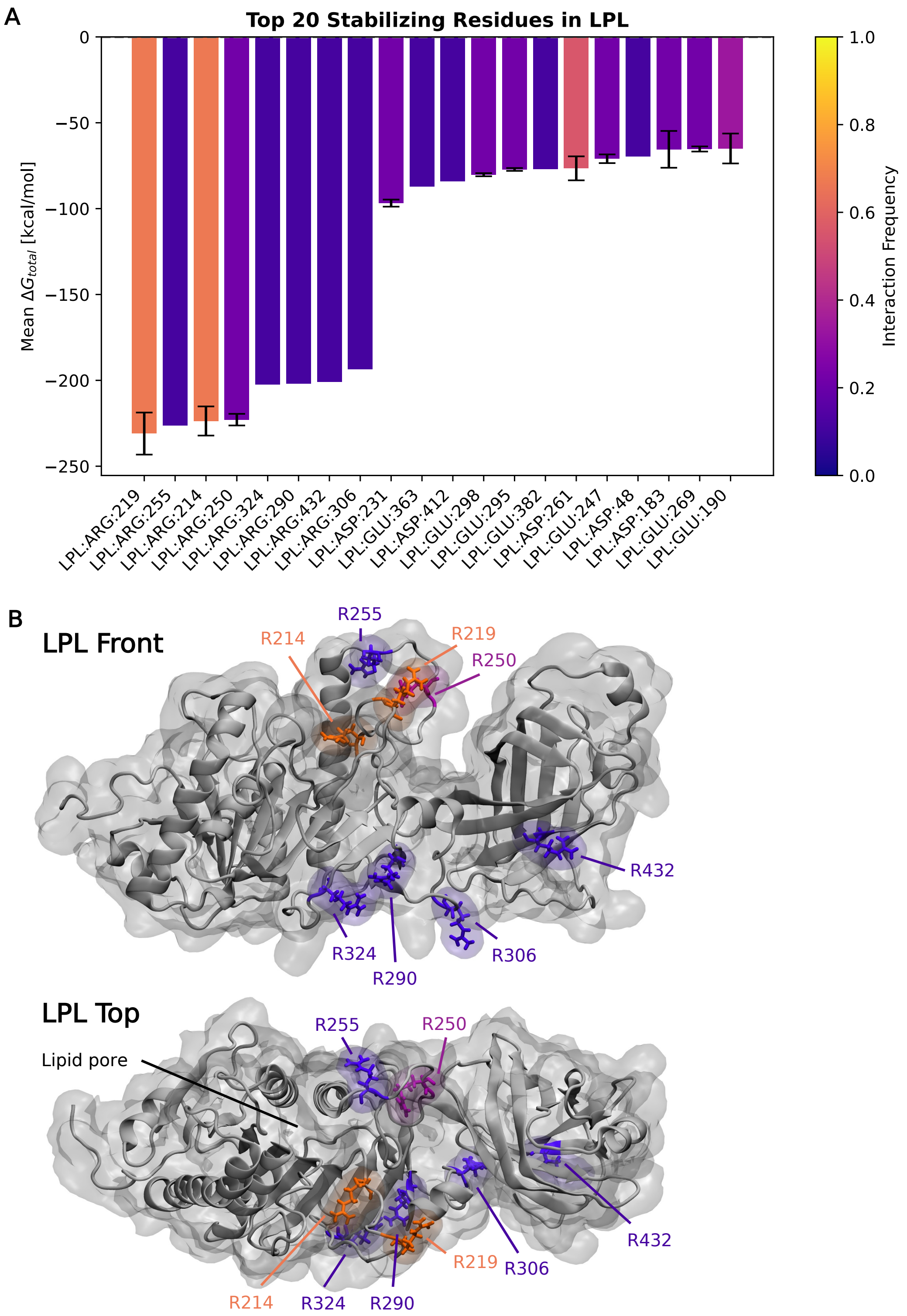


**Supplementary Figure 13. Stabilizing energy contributions from LPL residues in interactions with ApoC-II-P.** (A) Decomposition analysis from gmx_MMPBSA analysis, where the mean $\Delta G_{total}$ values for the top twenty stabilizing LPL residues are shown. The interaction frequency per residue is plotted as a heatmap as a fraction from 0-1. (B) Visualization of LPL, with the top eight residues colored according to their heatmap values in (A). Front and top views of LPL are shown.

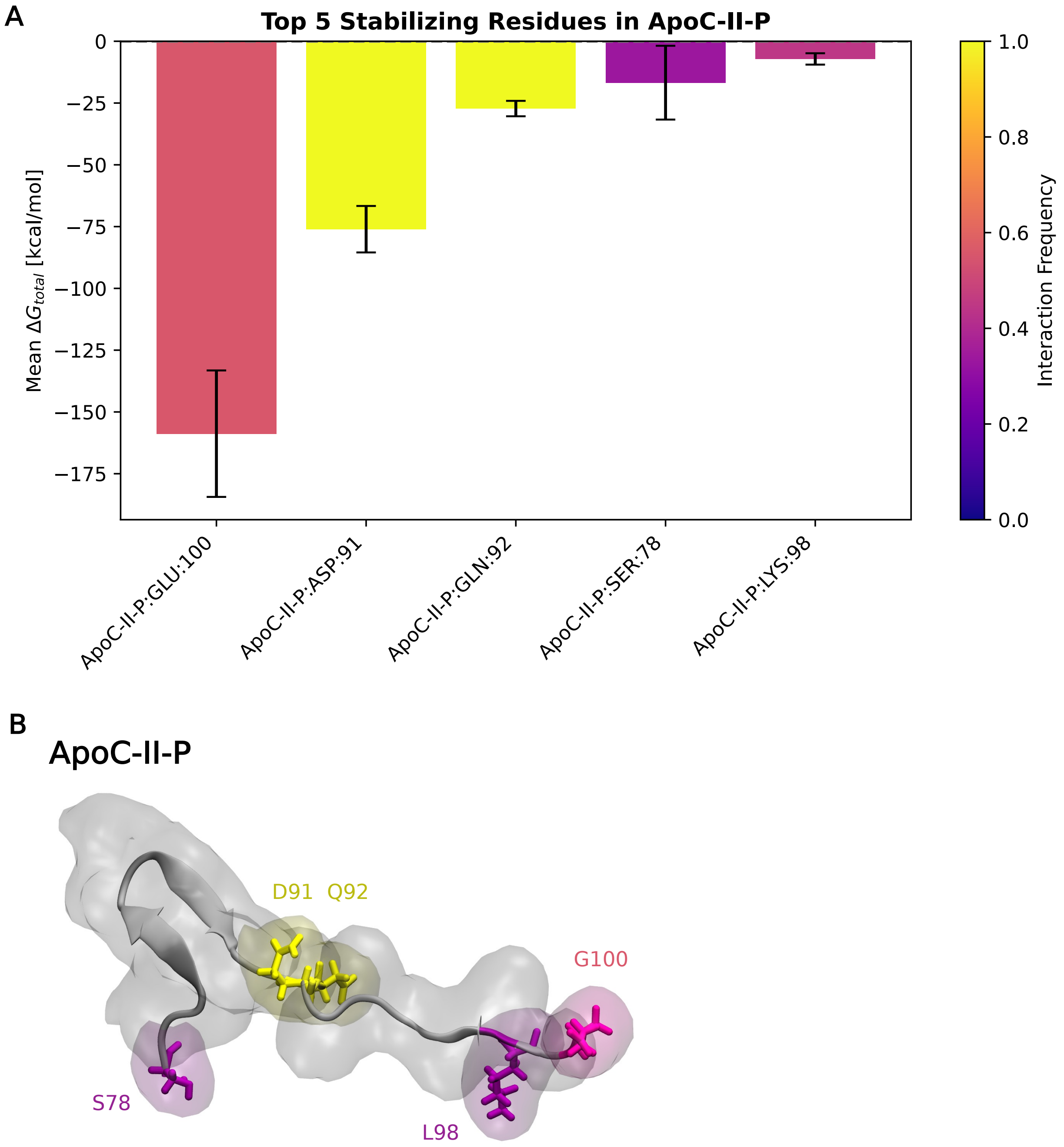


**Supplementary Figure 14. Stabilizing energy contributions from ApoC-II-P residues in interactions with LPL.** (A) Decomposition analysis from gmx_MMPBSA analysis, where the mean $\Delta G_{total}$ values for the top five stabilizing ApoC-II-P residues are shown. The interaction frequency per residue is plotted as a heatmap as a fraction from 0-1. (B) Visualization of ApoC-II-P, with the top five residues colored according to their heatmap values in (A).

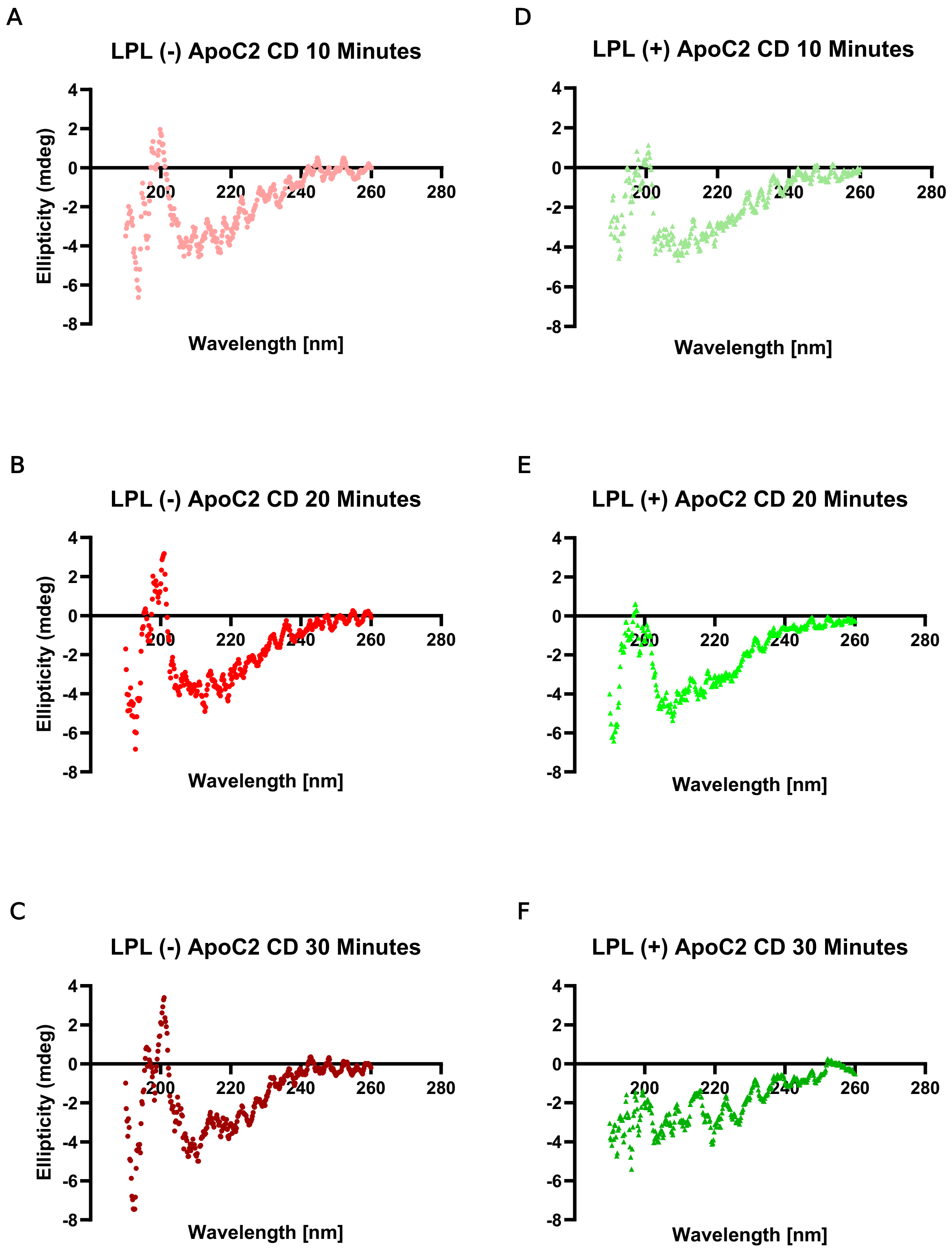


**Supplementary Figure 15. Circular dichroism (CD) spectra of bLPL with and without human ApoC-II-P, collected at 10-minute intervals over a 30-minute period**. (A–C) bLPL alone at 10, 20, and 30 minutes, respectively. (D–F) bLPL with human ApoC-II-P at the same time points.

**Supplementary Table 1. Number of data points (n) from the independent simulation runs (N) per group.** Values of n correspond to box plots used in Figs. 3E,6D and Supplementary Figs. 1A-C,7A-C. * corresponds to a sample group with <1% of trajectory sampling time across simulations and thus was not included in analyses.

| Group | n |
|---|---|
| Open (No ApoC-II-P) | 744 |
| Open Post-Flip (No ApoC-II-P) | 111 |
| Open (ApoC-II-P Far) | 486 |
| Open (ApoC-II-P Near) | 2828 |
| Open Post-Flip (ApoC-II-P Near) | 550 |
| Intermediate (No ApoC-II-P) | 3089 |
| Intermediate Post-Flip (No ApoC-II-P) | 794 |
| Intermediate (ApoC-II-P Far) | 667 |
| Intermediate (ApoC-II-P Near) | 10218 |
| Intermediate Post-Flip (ApoC-II-P Near) | 2693 |
| Closed (No ApoC-II-P) | 1265 |
| Closed (ApoC-II-P Far)* | 73 |
| Closed (ApoC-II-P Near) | 494 |

**Supplementary Table 2. LPL residues within 5 Å of ApoC-II-P in at least 50% of the converged, end time of the simulations (900-1000 ns).** ApoC-II binding sites are listed by the four numbered residue ranges along with N-terminal domain (NTD) residues outside of these domains and residues in the C-terminal domain (CTD).

| Number of Residues | | LPL Residues ≤5Å from ApoC-II-P from 900-1000 ns, for 50+% of Time Frame |
|---|---|---|
| **LPL X-R1** | | |
| aa77-87: | 0 | |
| aa114-126: | 0 | |
| aa212-244: | 0 | |
| aa245-263: | 0 | |
| NTD – other: | 5 | 292, 301, 304, 306, 307 |
| CTD: | 4 | 346, 432, 434, 441 |
| **LPL X-R2** | | |
| aa77-87: | 1 | 82 |
| aa114-126: | 3 | 120, 121, 122 |
| aa212-244: | 10 | 212, 213, 214, 215, 216, 217, 221, 237, 238, 239 |
| aa245-263: | 8 | 245, 246, 247, 255, 257, 258, 260, 261 |
| NTD – other: | 11 | 186, 187, 264, 268, 294, 295, 297, 298, 299, 301, 331 |
| CTD: | 14 | 344, 346, 379, 380, 381, 382, 387, 388, 389, 390, 391, 392, 393, 394 |
| **LPL X-R3** | | |
| aa77-87: | 0 | |
| aa114-126: | 0 | |
| aa212-244: | 17 | 212, 213, 214, 215, 216, 217, 218, 219, 221, 222, 223, 231, 233, 237, 238, 239, 244 |
| aa245-263: | 11 | 245, 246, 247, 248, 249, 250, 251, 252, 257, 260, 263 |
| NTD – other: | 8 | 190, 264, 292, 293, 294, 295, 327, 329 |
| CTD: | 12 | 361, 362, 363, 377, 378, 379, 410, 411, 412, 413, 414, 415 |
| **LPL Y-R1** | | |
| aa77-87: | 1 | 82 |
| aa114-126: | 1 | 121 |
| aa212-244: | 17 | 212, 213, 214, 215, 216, 217, 218, 219, 220, 221, 235, 236, 237, 238, 239, 240, 241 |
| aa245-263: | 8 | 249, 250, 252, 253, 255, 257, 260, 261 |
| NTD – other: | 6 | 187, 190, 191, 264, 268, 294 |
| CTD: | 0 | |
| **LPL Y-R2** | | |
| aa77-87: | 0 | |
| aa114-126: | 0 | |
| aa212-244: | 16 | 213, 214, 215, 216, 217, 218, 219, 220, 221, 222, 223, 224, 233, 239, 240, 241 |
| aa245-263: | 6 | 251, 252, 253, 257, 260, 263 |
| NTD – other: | 13 | 264, 290, 291, 292, 293, 294, 296, 301, 302, 304, 306, 307, 327 |
| CTD: | 0 | |
| **LPL Y-R3** | | |
| aa77-87: | 0 | |
| aa114-126: | 0 | |
| aa212-244: | 18 | 212, 213, 214, 215, 216, 217, 219, 220, 221, 222, 224, 233, 234, 235, 238, 239, 240, 241 |
| aa245-263: | 9 | 251, 252, 253, 254, 255, 256, 257, 260, 263 |
| NTD – other: | 5 | 190, 191, 210, 211, 264 |
| CTD: | 0 | |
| **LPL Z-R1** | | |
| aa77-87: | 5 | 81, 82, 83, 84, 85 |
| aa114-126: | 3 | 120, 121, 122 |
| aa212-244: | 4 | 219, 220, 221, 222 |
| aa245-263: | 1 | 261 |
| NTD – other: | 25 | 47, 48, 89, 90, 91, 93, 94, 158, 159, 160, 183, 184, 185, 186, 187, 188, 211, 264, 265, 266, 268, 269, 272, 273, 276 |
| CTD: | 0 | |
| **LPL Z-R2** | | |
| aa77-87: | 1 | 82 |
| aa114-126: | 3 | 120, 121, 122 |
| aa212-244: | 6 | 212, 214, 215, 216, 238, 239 |
| aa245-263: | 11 | 249, 250, 252, 253, 254, 256, 257, 258, 260, 261, 263 |
| NTD – other: | 6 | 185, 186, 187, 188, 264, 268 |
| CTD: | 0 | |
| **LPL Z-R3** | | |
| aa77-87: | 4 | 82, 83, 84, 85 |
| aa114-126: | 1 | 121 |
| aa212-244: | 9 | 212, 213, 214, 215, 219, 236, 237, 238, 239 |
| aa245-263: | 8 | 249, 252, 257, 258, 260, 261, 262, 263 |
| NTD – other: | 12 | 89, 91, 158, 159, 160, 185, 187, 264, 265, 268, 269, 294 |
| CTD: | 0 | |
